# Quantification of astrocytic synaptic pruning in hippocampus in response to *in vitro* Aβ oligomer incubation via colocalization analysis with C1q

**DOI:** 10.1101/2022.05.09.491159

**Authors:** Arpit Kumar Pradhan, Qinfang Shi, Katharina Johanna Tartler, Gerhard Rammes

## Abstract

Quantification of synaptic engulfment is an indirect measurement of synaptic pruning. Here, we provide a detailed protocol for the volumetric rendering of individual high-resolution astrocytes in the CA1 region of hippocampus in an *in vitro* slice model of Amyloid-beta (Aβ) treatment. This includes free floating slice preparation, treatment with Aβ oligomers, immunofluorescence, confocal imaging and analysis of individual astrocytes. We also provide a comprehensive analysis for 3D rendering of astrocytes and assessment of synaptic engulfment via “eat-me tag” C1q protein and synaptic marker PSD95.

**Highlights:**

- Measurement of synaptic engulfment in response to treatment with Aβ peptide
- Volumetric reconstruction of high resolution individual astrocyte
- Colocalization analysis of astrocyte and complementary “eat-me” protein C1q

**Graphical abstract:** 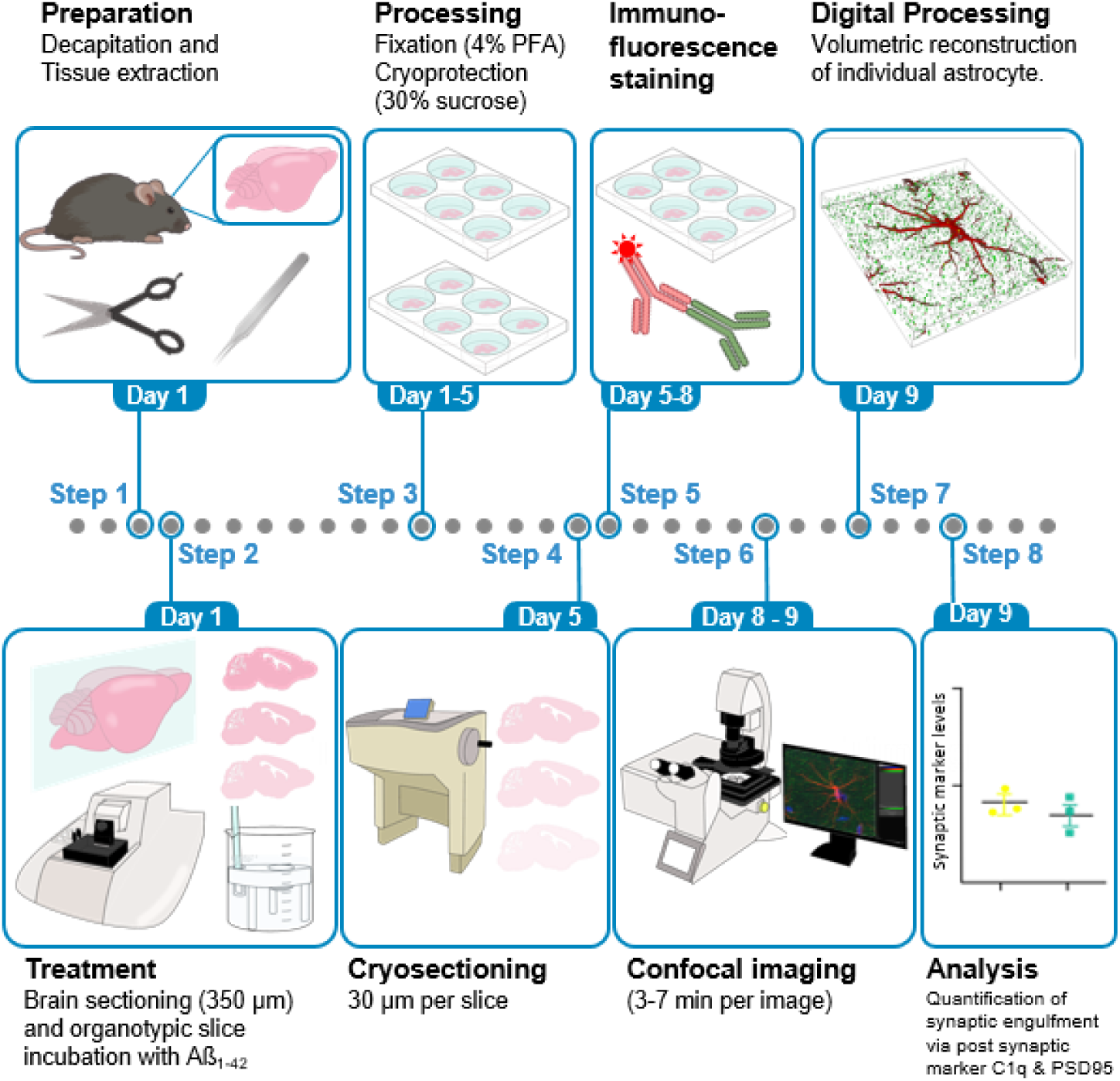

## Before you begin

Synaptic pruning, which is primarily mediated by the glial cells, is an essential step in modulation and shaping up of synapses and fine-tuning of neuronal connection (Tierney and Nelson III, 2009; Tau and Peterson, 2010; Perez-Catalan, Doe and Ackerman, 2021). Astrocytes, which form a bigger subset of glial cells, play a significant role in restructuring of synapses by regular crosstalk with microglia (Chung, Allen and Eroglu, 2015). This crosstalk is mediated by a diverse range of cytokines (Garland, Hartnell and Boche, 2022). C1q of the complement cascade, is one such essential tags which is also referred to as the “eat-me” signal and plays a critical role as a signaling molecule in the pruning of neurons (Iram *et al*., 2016) (Kovács *et al*., 2021). The pruning of synapses in the brain occurs via both C1q dependent and independent pathways (Györffy *et al*., 2018). Astrocytes upon sensing unnecessary synapses, release tumor growth factor-β (TGF- β), which increase the expression of C1q tags (Allen and Eroglu, 2017). Microglia upon recognizing this tag release inflammatory cytokine to engulf the synapse through phagocytosis. Previously, it has been reported that C1q expression is increased and associated with synapses before plaque deposition and is necessary for mediating the toxic effects of soluble Aβ oligomers on synapses and long-term potentiation (LTP) in hippocampus (Hong *et al*., 2016). From previous studies including one from our lab, the treatment of hippocampal slice with 50 nM Aβ_42_ caused a significant decline in the CA1-LTP of the hippocampus (Li *et al*., 2011; Rammes *et al*., 2018). The aim was therefore to look at the molecular level changes, particularly at the level of synapse, after Aβ_42_ incubation. We recently developed a protocol to analyze and systematically quantify synaptic engulfment of materials by the astrocytes in the hippocampus after *in vitro* slice treatment with Aβ_42_ oligomers. In order to look at the synaptic involvement, we also tried to look at the C1q engulfment in the astrocytic volume. Although this protocol has been optimized in the stratum radiatum layer of CA1 region of hippocampus, it can also be applied to other brain regions. This protocol of synaptic engulfment determination using *in vitro* slice treatment of Aβ_42_ oligomers can be combined with different treatment regimens of drugs/small molecules which affect the binding of Aβ_42_ oligomer.

### Preparation of preparation ringer and mess ringer solution [Part 1 and 2]

#### Timing: [30 min]

1. Prepare 1000ml of mess ringer (aCSF) solution
  a. Dissolve the following reagents in 800ml of milliQ water by constantly stirring the contents with a magnetic stirrer and bring up the level to 1000ml after the solution becomes transparent. Store the mess ringer solution in 4ºC for a maximum of seven days.

#### CRITICAL

The mess ringer solution should be freshly prepared before the starting of the week and should be immediately stored in 4ºC for a maximum of 7 days. Post this period the solution should not be used and another fresh batch should be prepared.

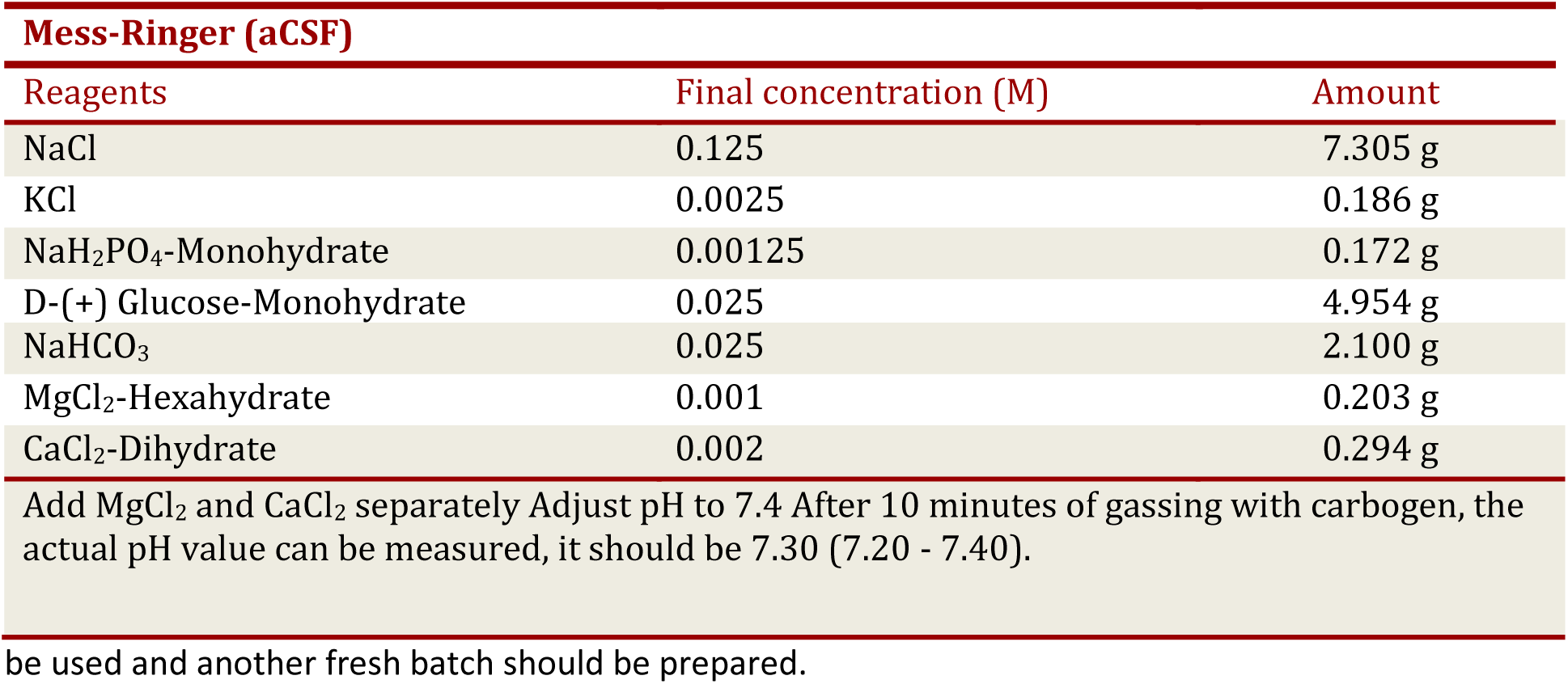

2. Prepare 2000 ml of preparation-ringer Solution
  a. Dissolve the following reagents in 1500ml of milliQ water by constant stirring with a magnetic stirrer and make up the volume to 2000ml.
  b. Divide the solution into 4 plastic bottles of 500ml each and store the bottles at -80ºC for a maximum of one month.

#### CRITICAL

The plastic bottles should not be completely filled, which would cause cracks in the bottle due to the expansion in volume. In an alternate scenario, where researchers intend to use the Preparation Ringer within a span of seven days, the 2000ml solution can directly be stored in the 4ºC and placed for 40 min in -80ºC before starting the sacrifice procedure.

| **Preparation-Ringer** |  |  |
| --- | --- | --- |
| Reagents | Final concentration (M) | Amount |
| NaCl | 0.125 | 14.610 g |
| KCl | 0.0025 | 0.373 g |
| NaH<sub>2</sub>PO<sub>4</sub>-Monohydrate | 0.00125 | 0.345 g |
| D-(+) Glucose-Monohydrate | 0.025 | 9.909 g |
| NaHCO<sub>3</sub> | 0.025 | 4.201 g |
| MgCl<sub>2</sub>-Hexahydrate | 0.006 | 2.440 g |
| CaCl<sub>2</sub>-Dihydrate | 0.00025 | 0.074 g |

Preparation Ringer within a span of seven days, the 2000ml solution can directly be stored in the 4°C and placed for 40 min in -80°C before starting the sacrifice procedure.

### Preparation of Amyloid-Beta (Aβ42) [Part 2]

#### Timing: [3-4 h]

1. 1 mg of Aβ_42_ is dissolved in hexafluoro-2-propanol (HFIP) (400 μl) and is incubated in room temperature (RT) until clear solution forms.

#### CRITICAL

This could take up to 90 minutes in room temperature. In an alternate scenario, one can warm up at 37ºC for 15-20 min.

2. Aliquot the solution into 20 tubes with 20 μl each.

#### CRITICAL

Use siliconized (low protein binding) tubes! Put the tubes directly on dry ice in a box.

3. Place the tubes in lyophilizer for up to 2 hours until white pellets form at the bottom of the tubes.

#### CRITICAL

Before placing the tubes into the lyophilizer, open them and close the tubes with parafilm while the lid is open. Gently poke several holes into the parafilm to make the lyophilisation process possible.

4. The tubes should be tightly closed, labelled and placed in -80ºC.
5. 100 μM stock of Aβ_42_ is made by using 111 μl DMSO. After adding DMSO, sonicate the tube on an ultrasonic bath for 15 minutes.

#### CRITICAL

Use only freshly opened DMSO. The stock solution can be kept in -20ºC and should not be kept out in the room temperature longer than 20-30 minutes.

### Preparation for slice incubation [Part 2]

#### Timing: [15 min]

1. Before sacrificing the mice, make sure that the microtome setup is working and running. The water bath should be initially switched on and kept at 35ºC.
2. Prepare two beakers (control and treatment) with 70ml of mess ringer solution and fumigate with carbogen (95% O_2_ and 5% CO_2_).

#### CRITICAL

Make sure that the glass filter is properly inserted inside the beaker containing mess ringer. If it is inserted too deep, there would be bubbles on the net which hampers the health of the slices. Additionally, make sure that the glass filter is not on the surface, which would lead to unequal distribution of carbogen. Ideally the glass filter should be inserted in a way in which it is placed at three-fourth the position of the stand (Figure 1).

3. Prepare a petri dish with mess ringer solution and fumigate it with carbogen.
4. The microtome box should be surrounded by ice all the time (this is to keep the preparation ringer solution in slushy-state for smooth cutting of the slices). Inside the microtome box fill in the preparation ringer solution (around 6 tablespoon) and start fumigating it with carbogen.

**Figure 1:**
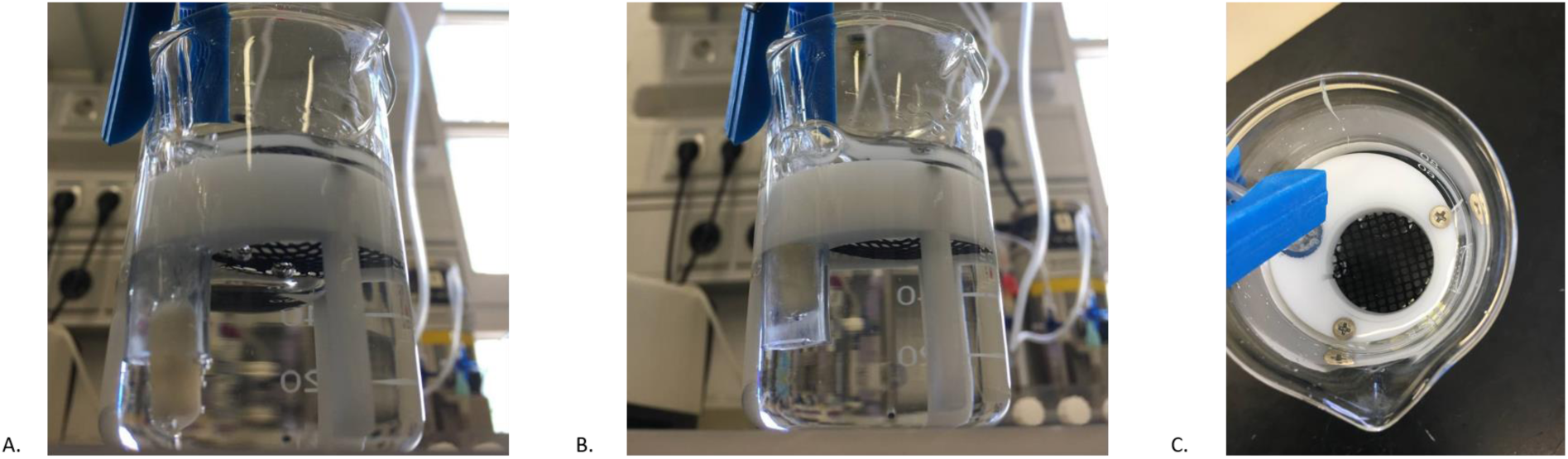
Positioning of micro filter candle for carbogen fumigation. A. Representative image of wrong positioning of the tube inside the beaker. Placing the tube completely inside the beaker will result in air bubble formation on the net which affects the health of the slice. B. Correct positioning of the tube inside the beaker. C. Air bubble formation in the nets should always be avoided.

#### CRITICAL

The preparation ringer should be always in a slushy state. It should not be completely in a liquid state or in a hard ice state. From our experience, we recommend placing a preparation ringer bottle kept initially in -80ºC, in 4ºC refrigerator a day before starting the experiment. In an alternate scenario, if the preparation ringer is kept in 4ºC, 500ml of it can be separated and kept in -80ºC for 40 min to bring it into a slushy state.

#### CRITICAL

Don’t place the razor inside the microtome yet! However, the razor can be placed inside the holder and can be kept in a safe place before starting the cutting.

### Preparation of buffer solution for immunofluorescence

#### Timing: [2 h]

1. Prepare 10x phosphate buffer saline (PBS). Dilute it 10 times to obtain 1X PBS. This can be stored in the room temperature for up to 6 months.
2. Prepare 4% paraformaldehyde (PFA) in 1x PBS and store it in 4ºC for a maximum of 7 days. It is recommended to start a fresh preparation of PFA before starting the staining procedure.
  a. Dissolve 20g of PFA powder in 500 ml of 1xPBS solution on a hot plate heated up to 50ºC and stirred with a magnetic stirrer continuously until the powder dissolves completely. After dissolving, adjust the pH value to 7.4.

#### CRITICAL

PFA is a hazardous chemical. The solution should be prepared inside a fuming hood.

3. Prepare 30% sucrose solution (for cryoprotection) in 1xPBS solution.
  a. Dissolve 150g of sucrose powder in 500 ml 1xPBS solution and mix the contents with magnetic stirrer.
4. Preparation of blocking solution (10% normal goat serum)
  a. Dissolve 1 ml of 100% normal goat serum (NGS) in 9 ml of 1xPBS + 0.3% triton-X 100 solution.

#### CRITICAL

The blocking solution should always be prepared freshly before every staining and should not be stored for a longer time.

## Key resources table

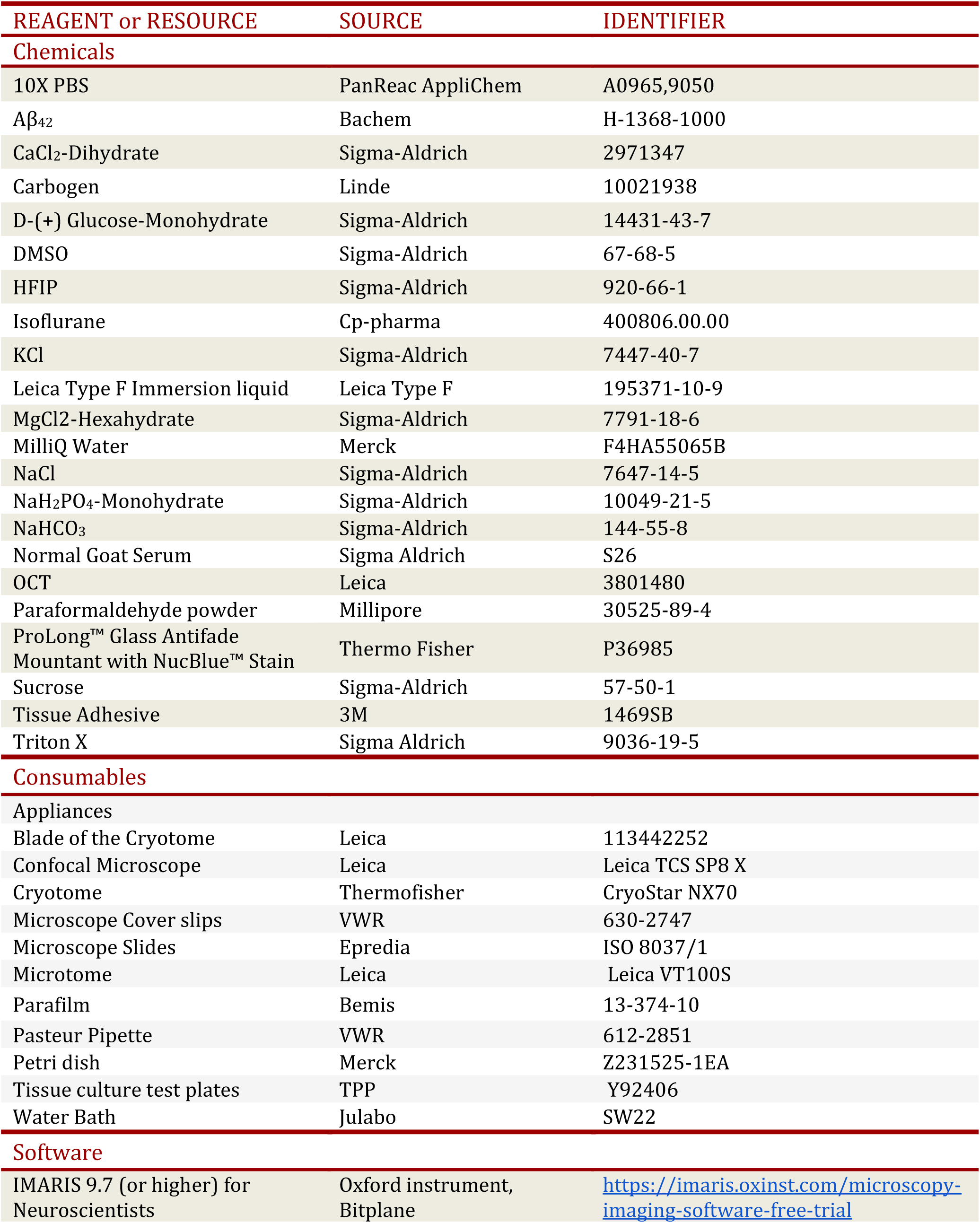
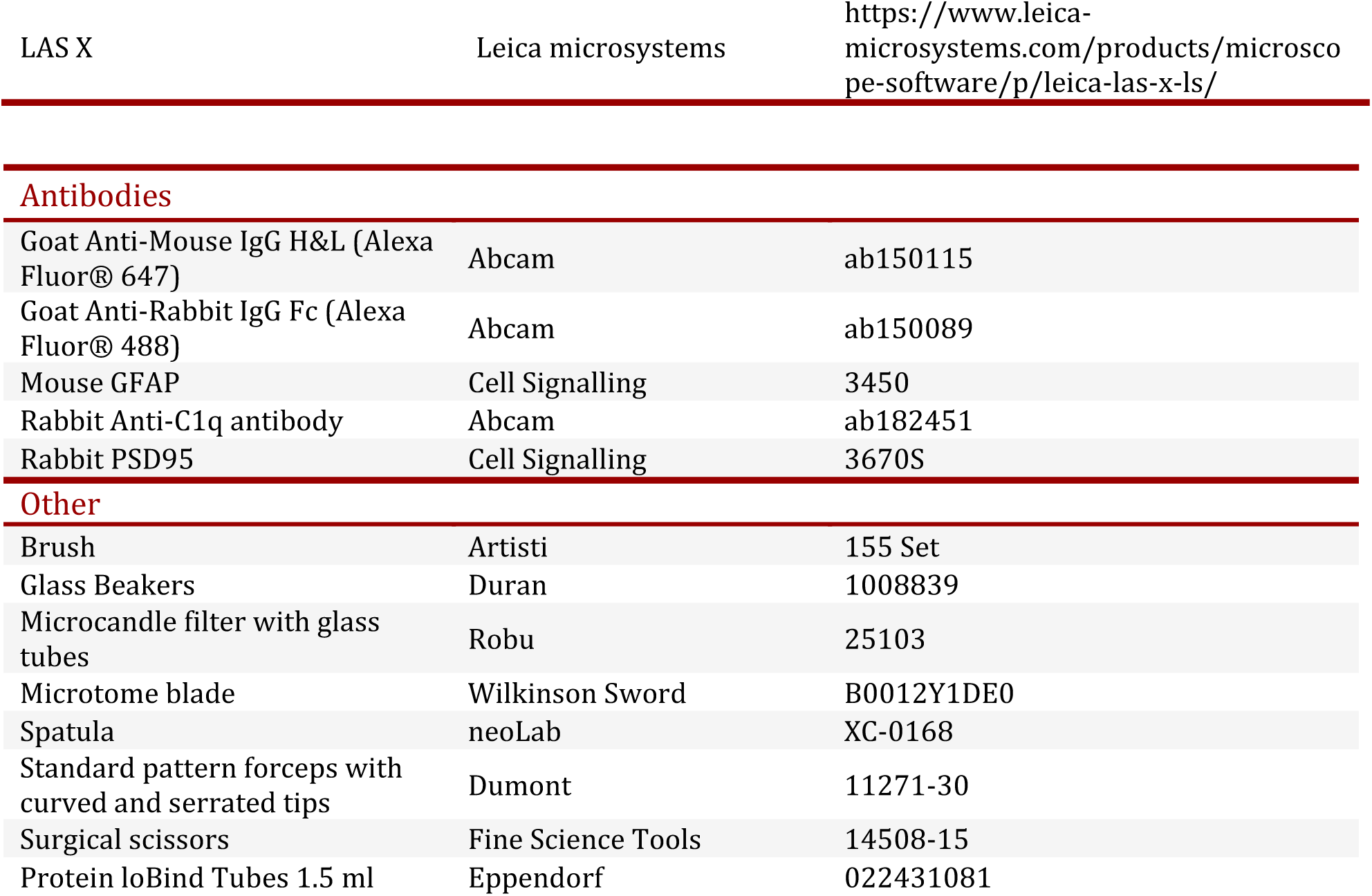

| REAGENT or RESOURCE | SOURCE | IDENTIFIER |
| --- | --- | --- |
| **Chemicals** |  |  |
| 10X PBS | PanReac AppliChem | A0965,9050 |
| Aβ<sub>42</sub> | Bachem | H-1368-1000 |
| CaCl<sub>2</sub>-Dihydrate | Sigma-Aldrich | 2971347 |
| Carbogen | Linde | 10021938 |
| D-(+) Glucose-Monohydrate | Sigma-Aldrich | 14431-43-7 |
| DMSO | Sigma-Aldrich | 67-68-5 |
| HFIP | Sigma-Aldrich | 920-66-1 |
| Isoflurane | Cp-pharma | 400806.00.00 |
| KCl | Sigma-Aldrich | 7447-40-7 |
| Leica Type F Immersion liquid | Leica Type F | 195371-10-9 |
| MgCl<sub>2</sub>-Hexahydrate | Sigma-Aldrich | 7791-18-6 |
| MilliQ Water | Merck | F4HA55065B |
| NaCl | Sigma-Aldrich | 7647-14-5 |
| NaH<sub>2</sub>PO<sub>4</sub>-Monohydrate | Sigma-Aldrich | 10049-21-5 |
| NaHCO<sub>3</sub> | Sigma-Aldrich | 144-55-8 |
| Normal Goat Serum | Sigma Aldrich | S26 |
| OCT | Leica | 3801480 |
| Paraformaldehyde powder | Millipore | 30525-89-4 |
| ProLong™ Glass Antifade Mountant with NucBlue™ Stain | Thermo Fisher | P36985 |
| Sucrose | Sigma-Aldrich | 57-50-1 |
| Tissue Adhesive | 3M | 1469SB |
| Triton X | Sigma Aldrich | 9036-19-5 |
| **Consumables** |  |  |
| Appliances |  |  |
| Blade of the Cryotome | Leica | 113442252 |
| Confocal Microscope | Leica | Leica TCS SP8 X |
| Cryotome | Thermofisher | CryoStar NX70 |
| Microscope Cover slips | VWR | 630-2747 |
| Microscope Slides | Epredia | ISO 8037/1 |
| Microtome | Leica | Leica VT100S |
| Parafilm | Bemis | 13-374-10 |
| Pasteur Pipette | VWR | 612-2851 |
| Petri dish | Merck | Z231525-1EA |
| Tissue culture test plates | TPP | Y92406 |
| Water Bath | Julabo | SW22 |
| **Software** |  |  |
| IMARIS 9.7 (or higher) for Neuroscientists | Oxford instrument, Bitplane | https://imaris.oxinst.com/microscopy-imaging-software-free-trial |
| Antibodies |  |  |
| Goat Anti-Mouse IgG H&L (Alexa Fluor® 647) | Abcam | ab150115 |
| Goat Anti-Rabbit IgG Fc (Alexa Fluor® 488) | Abcam | ab150089 |
| Mouse GFAP | Cell Signalling | 3450 |
| Rabbit Anti-C1q antibody | Abcam | ab182451 |
| Rabbit PSD95 | Cell Signalling | 3670S |
| Other |  |  |
| Brush | Artisti | 155 Set |
| Glass Beakers | Duran | 1008839 |
| Microcandle filter with glass tubes | Robu | 25103 |
| Microtome blade | Wilkinson Sword | B0012Y1DE0 |
| Spatula | neoLab | XC-0168 |
| Standard pattern forceps with curved and serrated tips | Dumont | 11271-30 |
| Surgical scissors | Fine Science Tools | 14508-15 |
| Protein loBind Tubes 1.5 ml | Eppendorf | 022431081 |

## Step-by-step method details

### Preparation: Decapitation and Tissue Extraction

#### Timing: 20 min

In the first step, the mice brain is decapitated and placed inside the preparation ringer solution which is constantly fumigated with the carbogen. The preparation of microtome and carbogen is to be done prior to sacrificing the animals (as described in “Before you begin” section). It should be noted that in order to keep the slices healthy for a longer duration, the carbogen should be constantly mixed with the respective preparation ringer or mess ringer solution.

1. Prepare the surgery site with surgical scissors, standard pattern forceps with curved and serrated tips, fresh blades and a spatula (Figure 2 A, B).
2. Keep a beaker with the preparation ringer solution continuously fumigated with carbogen at the side. Make sure to keep the beaker in a box full of ice, to keep the preparation ringer in a slushy state. Additionally, keep another petri dish with preparation ringer supplied with carbogen, which serves as a platform to take off the brain from the decapitated head.
3. The mice is anesthetized by placing it in an isoflurane chamber with 5% isoflurane concentration and at an oxygen flow rate of about 2L/min. Wait till the animal is anesthetized. The standard and a reliable way to test whether the mice is aptly anesthetized is by doing a twofold check. The first one is to check for the instance when the mice loses the righting reflex. The second one is by pinching the tails with the forceps. Once, checked for these measures, the mice can be brought to the guillotine and the head is decapitated.

**Figure 2:**
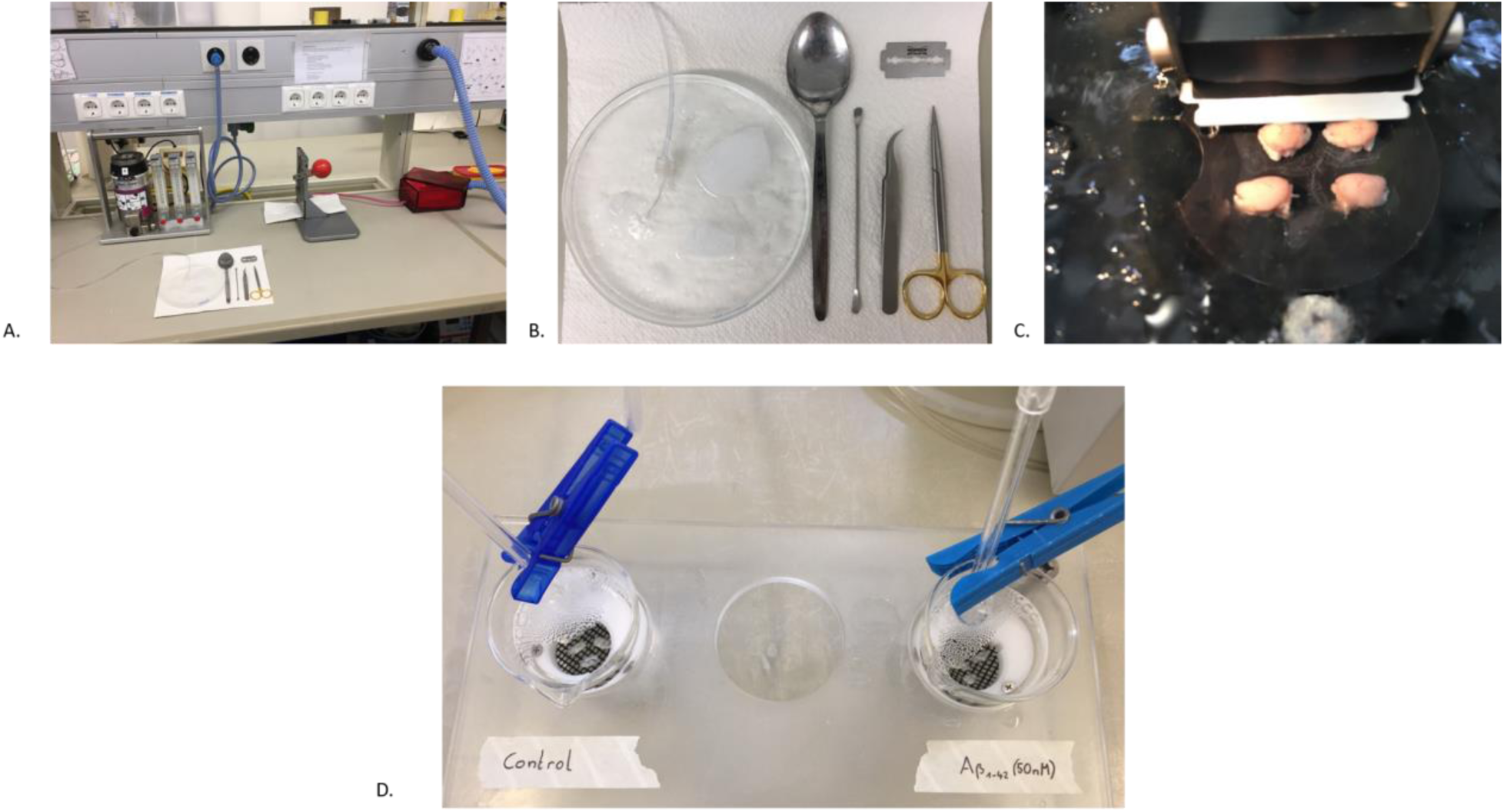
Preparation of brain slices and incubation with Aβ _42_ peptide. A. Overview of the setup required for the sacrifice of the mice. Isoflurane (5% Concentration) was used as the anesthetizing agent. B. Close overview of the surgical set required for the dissection of mice. From the left, spoon, spatula, standard pattern forceps with curved tips and surgical scissor. C. Microtome cutting through the sagittal section of the mouse brain. D. Organotypic incubation of slice with Aβ_42_ peptide.

**Note**: It is advised to use younger mice of 6-8 weeks for this method of *in vitro* slice treatment with Aβ oligomers.

4. Quickly transfer the decapitated head to the petri dish containing preparation ringer solution. With the help of sharp surgical scissors, cut open the scalp through the midline along the sagittal suture. Make a lateral cut on both the sides of skull and remove the top of the skull to expose the brain.

#### CRITICAL

Be careful while cutting through the joints of the skull particularly the bregma. If the bone doesn’t fall off completely, it can be slowly removed using a fine-ended forceps.

5. From the caudal side, gently lift out the brain using the spatula. Quickly transfer the brain to cooled preparation ringer solution in the beaker. Discard mouse body and head in the freezer, disinfect and clean guillotine, anesthetizing chamber and workspace.
6. Bring the brain closer to the microtome setup.

### Treatment: Sectioning (350 um per slice) and incubation with Aβ42

#### Timing: 5 h

The brain is sectioned in the sagittal plane to 350 um slices for incubation and treatment with the Aβ oligomers. Sagittal section offers a good frame of view for the stratum radiatum layer of CA1 region of hippocampus. This phase includes the incubation of slice for 30 min in water bath at 35ºC and then in the aCSF at room temperature for 1 hour. This allows sufficient time for the recovery of slices.

7. Transfer the brain to a small flat platform and cut off the cerebellum. Separate the two hemisphere with a sharp razor. Put two streak of tissue adhesive on the microtome cutting plate.
8. Dry both the hemispheres of the brain by first transferring it on a filter paper for a short while. Transfer the brain hemispheres to the glue on the microtome plate and place the plate inside the holder. Carefully lock the plate and insert the razor inside the microtome.
9. Fix the cutting window in the microtome by setting the starting and the final position of brain. Set the thickness to 350 um. Start trimming the hemisphere until anterior hippocampus starts to appear (Figure 2 C). Start collecting the slices and divide them into the respective beaker: control and treatment (Aβ_42_). Cut slices and carefully transfer them to fumigated mess-ringer filled beakers in the water bath with a pipette (Figure 2 D).

#### CRITICAL

Always be watchful of the bubbles in the nets holding the slices. They directly affect the longevity of the slices. Also make sure that the brain slices are not overlapping on each other. This would affect the health of the slices.

#### CRITICAL

In order to reduce the overcrowding of slices inside the beaker, one can trim off the rest of the brain area from the sagittal section of the brain keeping the hippocampus and cortex intact. This allows more slices to be accommodated in to the respective beakers.

10. After completing the sectioning, place the beakers containing the slices in the water bath at 35ºC for about 30 min and then remove it from the water bath and place it in the room temperature for 1 hour. This allows sufficient time for the revival of slice.
11. Follow the incubation scheme as shown in the schematics. For 70 ml of the aCSF solution, pipette out 70 μl of Aβ_42_ (50 nM), into the treatment beaker and incubate it for a duration of 90 mins. The Aβ_42_ would evenly mix by the continuous fumigation of carbogen. The slices in the control chamber can be left in the aCSF with continuous supply of carbogen for the same 90 min duration.

|  |  |  |  |
| --- | --- | --- | --- |
| Control | 30 min at $35^{\circ}\text{C}$ | 1 h at RT in aCSF | 1.5 h |
| $\text{A}\beta_{1-42}$<br>Treatment | 30 min at $35^{\circ}\text{C}$ | 1 h at RT in aCSF | $\text{A}\beta_{1-42}$ (50 nM)<br>1.5 h |

#### CRITICAL

Do not keep the Aβ_42_ stock solution for longer time at room temperature. 20 minutes prior to the starting of incubation with the amyloid beta oligomer, the Aβ_42_ can be brought to the room temperature to allow the contents to thaw.

### Processing of brain slices

#### Timing: 3 days

This step is crucial for fixation and further cryopreservation. Hard Fixatives like PFA allow covalent cross-linking between molecules and stabilizes the protein. The slice are then placed in 30% sucrose solution for cryoprotection.

12. Divide the slices into a 6 well plate and pour in adequate amount of 4% PFA to cover the slices. The slices should be incubated overnight on a 3D shaker at 4ºC with PFA.

#### CRITICAL

PFA is hazardous. Take adequate care while handling it. It should be freshly prepared and should not be reused for multiple times.

13. On the following day, wash the slice three times with 1xPBS for 10 minutes each.
14. Pippete out 2 ml of 30% sucrose solution into each well and incubate it for 2 days on 3D shaker at 4ºC.

### Cryosectioning: 30 um per brain slice

#### Timing: 2-3 h per brain

The slices are further sectioned to 30 μm of thickness for better staining and effective immunofluorescence protocol. This step is crucial for obtaining good free-floating slices. Cracks or tear in the slices would directly affect the staining and mounting onto slide.

15. Pour in a small amount of the OCT media on the sample disk and allow it to solidify. Carefully place the disk in the holder and trim the OCT compound until it becomes a flat surface.

#### CRITICAL

Make sure that the surface of the OCT is completely flat. The brain slices (350 μm) when kept on an uneven surface, would result in unequal thickness of slices and would directly affect the staining process.

16. Take the specimen disk out, mark the position where you intend to keep the brain slice. For our purpose, we marked the OCT with position resembling the hands of a clock. The brain slice is carefully picked up by a spatula and is placed horizontally on the specimen desk. Now cover the surface of slice with ample amount of OCT media and place the specimen disk back in the holder. Start trimming until the slices are reached. Once the slices are reached, set to desired thickness (30 μm) (Figure 3).

**Figure 3:**
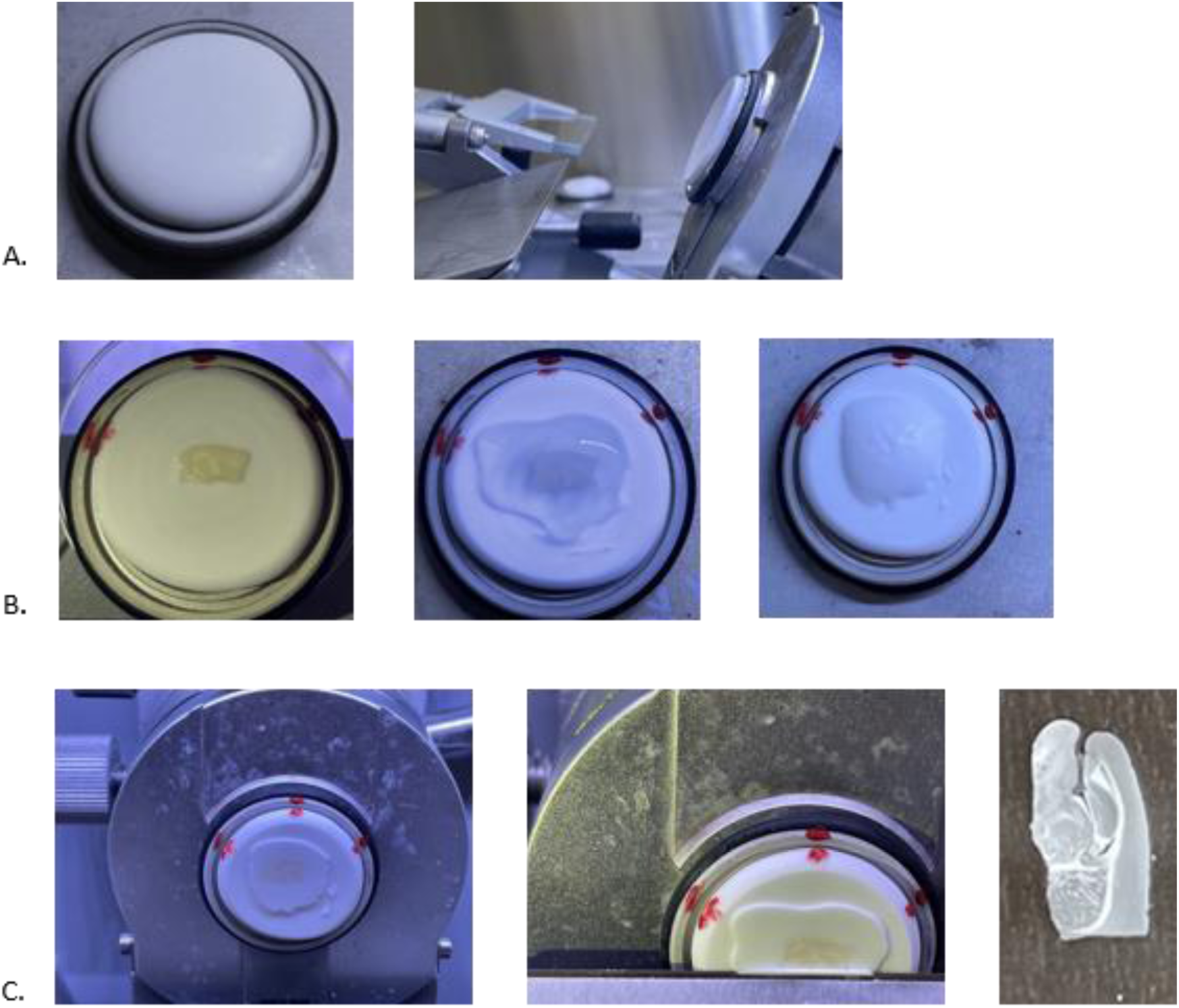
Cryosectioning of brain slices (30 μm): A. The first step is to cover the specimen disk with the OCT mounting media. Flatten the surface by initially trimming off the OCT media and creating a smooth flat surface. B. Place the brain slice (350 μm) horizontally on the surface of flattened OCT mounting medium. If not flat, gently press the brain slice to the surface and cover the surface with OCT C. Start trimming of extra parts before reaching the brain slice. Thin straight 30 μm sections as seen in the last image of the panel is produced at the end of this procedure.

#### CRITICAL

It is essential to make sure that the slice are laid flat on the OCT media. If necessary, it can softly be pressed using a brush to make the surface flat.

17. Carefully cut 3-4 slices, take them up all at once with a metal pick and transfer them into six-well plate filled with 1x PBS (the slices will unroll in there).

#### CRITICAL

In an alternate scenario, if one wishes to do the staining not in a free-floating slice, rather on a slide, the brain slice can be made flat using an anti-roll plate and can directly be collected using the slide. These slides can be stored in -80ºC for a longer duration.

18. It is possible to obtain 6-7 good 30 μm slices from a single 350 μm sagittal sectioned hippocampus slice. Collect enough slices in 1xPBS from both control as well as treatment group.

### Immunofluorescence Staining

#### Timing: 3 days

In order to assess the synaptic engulfment of the “eat me signal” C1q protein, we perform the immunofluorescence on a sagittal section. The synaptic engulfment is further verified using a post-synaptic marker PSD95. The astrocytes are marked by using GFAP. Pre-select good slices using a brush and place them in the 6 well plate. Avoid putting a large number of slices into a single well, which would lead to overcrowding. A maximum of 3-4 slices per well is recommended.

19. Wash the slices 3 times with 1x PBS using a plastic Pasteur pipette. Handle the slices carefully.
20. Blocking: Pipette out 1ml/well of blocking solution 10 % NGS in 1xPBS+0.3 % triton-X and block for a duration of 2 hours in room temperature on a 3D shaker.
21. Remove the blocking solution. Incubate with primary antibody (800 μl/well, diluted in 10% NGS) for 2 days at 4ºC. Rabbit Anti C1q (1:250) for staining C1q protein, mouse anti GFAP (1:800) for staining astrocytes and rabbit anti-PSD95 (1:400) were used.
22. Remove the primary antibody and put it in a separate tube marked as recycled. These reused antibodies can be used for another round of staining.
23. Wash the slice 3 times with 1X PBS solution for 10 min each. Meanwhile prepare the secondary antibody.
24. Incubate the slices with respective secondary antibodies anti-mouse 647 (1:500) and anti-rabbit 488 for 2 hours at room temperature. Cover the plate with a shielded cover (like aluminum foil) to keep it dark.
25. Wash the slice 3 times with 1xPBS solution for 10 min each.
26. Take a petri dish filled with PBS and transfer few slice from the 6 well plate into petri dish and carefully place them flat on the slide. Use a brush to transfer and flatten them.

**NOTE**: For an efficient and easy transfer of the slice onto the slides, keep the slide tilted with half of its surface inside PBS. Carefully move up the slice in the junction where the slide touches PBS and slowly pull out the slide. The slice sticks onto the surface. If there are still small folds on the slice, use a soft fine-ended brush and carefully adjust the slice to make it flat.

27. Keep the slides tilted on filter paper in dark to dry. Mount the slices with a DAPI mounting media.

#### CRITICAL

Each slide should not contain more than 4 brain sections. Extreme care should be taken while putting coverslip on the slice. Put adequate amount of the mounting media to avoid putting pressure on the slice which could affect the structure of the proteins under study and thereby impacting rendering process in the further steps. Also make sure not to introduce any air bubble into the specimen. An easier way to accomplish this would be to put a thin pipette head between the coverslip and the slide and slowly removing the pipette allowing the mounting media to slowly but evenly distribute above the sample. In any case, if there is still air bubble, avoid scratching the coverslip or applying direct pressure from the top, instead start removing the bubble by slowly applying pressure from the sides, particularly from the areas which are distant from the brain specimen.

#### CRITICAL

For preserving the fluorescent signals for a longer period of time, keep the slices in -20ºC. When kept in the room temperature, the fluorescent signal decays. Before imaging the slices on a confocal scope, they can be briefly kept in the room temperature (20-30 min).

### Confocal Imaging: “Hunting for Astrocytes”

#### Timing: 2 h per brain

We image high resolution individual astrocytes from the stratum radiatum layer of CA1 region of hippocampus. The slides are double-immunofluorescent stained having astrocyte and either the complementary tag C1q protein or a postsynaptic marker PSD95 for reference. The images taken during the process are further deconvoluted by using the lightening function in the Leica confocal scope. This is to obtain a better signal to noise ratio. For effective volumetric analysis of the synaptic engulfment by the astrocyte, it is essential to reduce the background noise. To effectively categorize the astrocyte, the slides were marked with the DAPI mounting media (nuclear stain). Since we are particularly focused on the CA1 area of hippocampus, it is also crucial to mark distinct layers and regions of the hippocampus, to effectively identify our region of interest and pick the astrocyte from the layer. At the time of imaging we require three active fluorescent channels. Leica Microscope SP8 provides the user an advantage to have 2 hybrid detector (HyD) which produces less noise. Apart from these HyD sensors, they also have an additional photomultiplier tube (PMT) sensor. Since the Leica microscope, has a maximum of 2 HyD sensors and the rest PMT sensors, it is essential to pick the correct proteins for the respective sensor. To make things easier, it is always advisable to keep large, ubiquitous or uniformly expressed proteins in the PMT sensor and the proteins of interest for colocalization and for the analysis of the synaptic engulfment to be kept in the HyD sensor. In our case, we put DAPI under the PMT sensor and GFAP and C1q/PSD95 in HyD sensor. Following were the settings of the microscope used for our acquisition of astrocytes (The settings, however, could vary greatly upon the region of interest, proteins for colocalization and the staining protocol):

**Imaging Setup:** Leica Confocal SP8 with lightening

**Frame size**: 1024 X 1024 pixels

**Pinhole**: 1 AU

**Lasers**: 499 nm (C1q, PSD95 Intensity: 5-10%), 405 nm (DAPI) and 653 nm (GFAP), laser power depends on staining efficacy

**Gain (Master, analog)**: 800 - 1000 V

**Digital Offset**: 0 for 405 (PMT), (HyD doesn’t require a digital offset)

**Scan Area**: Zoom according to the focused astrocyte

**Averaging**: Averaging 2 line and displaying the mean

**Bit Depth**: 8-bit, bidirectional scanning

**Scanning speed**: 200 Hz

**Z-stack scanning**: System optimized (normally 0.3 mm), scanning from the top to bottom of the focused astrocyte.

**Objective**: 63 oil (Leica Type F Immersion liquid n_e23_ = 1,5180, v_e_ = 46).

**Selection of channels**: GFAP (Alexa 647, Cell Signaling, red, 653 nm laser)

C1q (Alexa 488, Abcam, green, 499 nm laser) PSD95 (Alex 488, Cell Signaling, 499 nm laser)

**Processing**: Lightning Function: Adaptive

Refractive Index: 1.44

(Parameter Settings depend on image quality/staining intensity, background etc.)

**Mounting Medium**: ProLong TM Glass Antifade Mountant with NucBlueTM Stain

28. The confocal should be switched on 30 minutes prior to starting the experiment. Care should be taken to calibrate the platform before imaging. Non-calibration will interfere while taking the z-stacks. The laser power should also be stabilized before imaging.
29. The primary step here is to locate the CA1 area of the hippocampus. In order to achieve this, start from a lower magnification (10x). Having DAPI in the regiment of fluorophores greatly helps in identifying the region of interest (CA1) (Figure 4).

**Figure 4:**
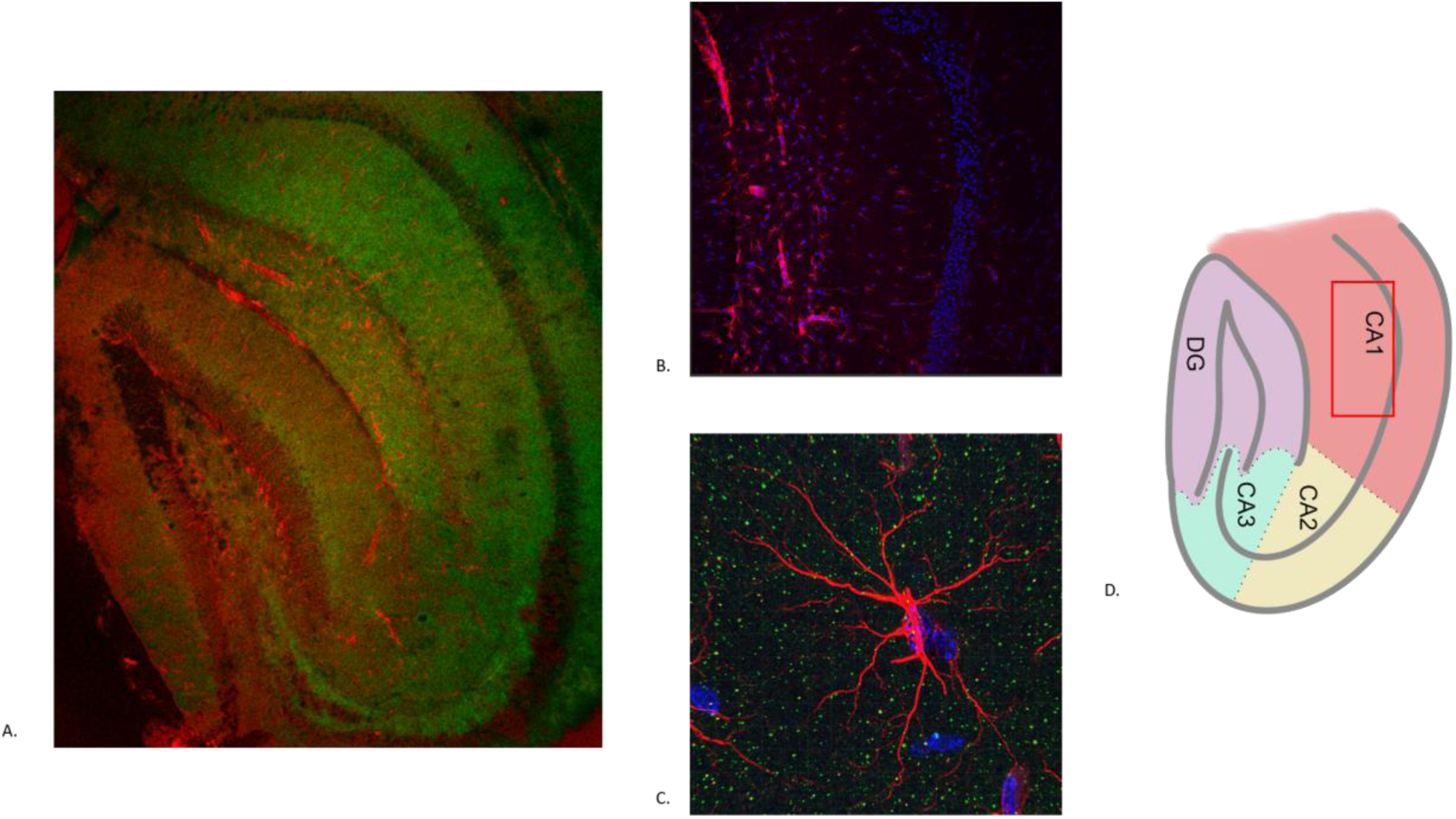
Overview of the C1q-GFAP interaction in the hippocampus. A. 10X magnification of the sagittal hippocampal slice detailing through the different region of hippocampus stained with C1q (green) and GFAP (red). B. 20X magnification focusing on the CA1 region of the hippocampus stained with DAPI (Blue) and astrocyte (Red). Just below the densely stained area (pyramidal layer) is the stratum radiatum layer where we pick the individual astrocyte. C. 63X magnification of a high resolution individual astrocyte (red) stained alongside with C1q (green) and DAPI (Blue). D. Schematic representation of different regions of hippocampus. The astrocytes are selected from the stratum radiatum layer which is illustrated in magenta and is marked using a red rectangular box.

**NOTE**: In our experience, it is easier to locate the dentate gyrus (V shaped structure) in the hippocampus first and then track down the CA1 area of the hippocampus. While finding the stratum radiatum of the CA1 can be tricky, it can eased by locating the pyramidal layer which is densely packed with nuclear body as seen in the DAPI channel. The stratum radiatum layer is directly below the pyramidal layer.

30. Once the region has been identified, switch to 63x (oil) for single cell acquisition of astrocytes. Do not put excess of the immersion oil.
31. Adjust all the three channels: Blue, Green and Red with the parameters as suggested above. Slowly move across the XY plane and look for well processed and structured astrocytes. The z-stack was set according to the GFAP channel in a way that it covers the entire volume of the astrocyte. The step size can be changed as per needs and demands of the experiment (Figure 5 A, B).

**Figure 5:**
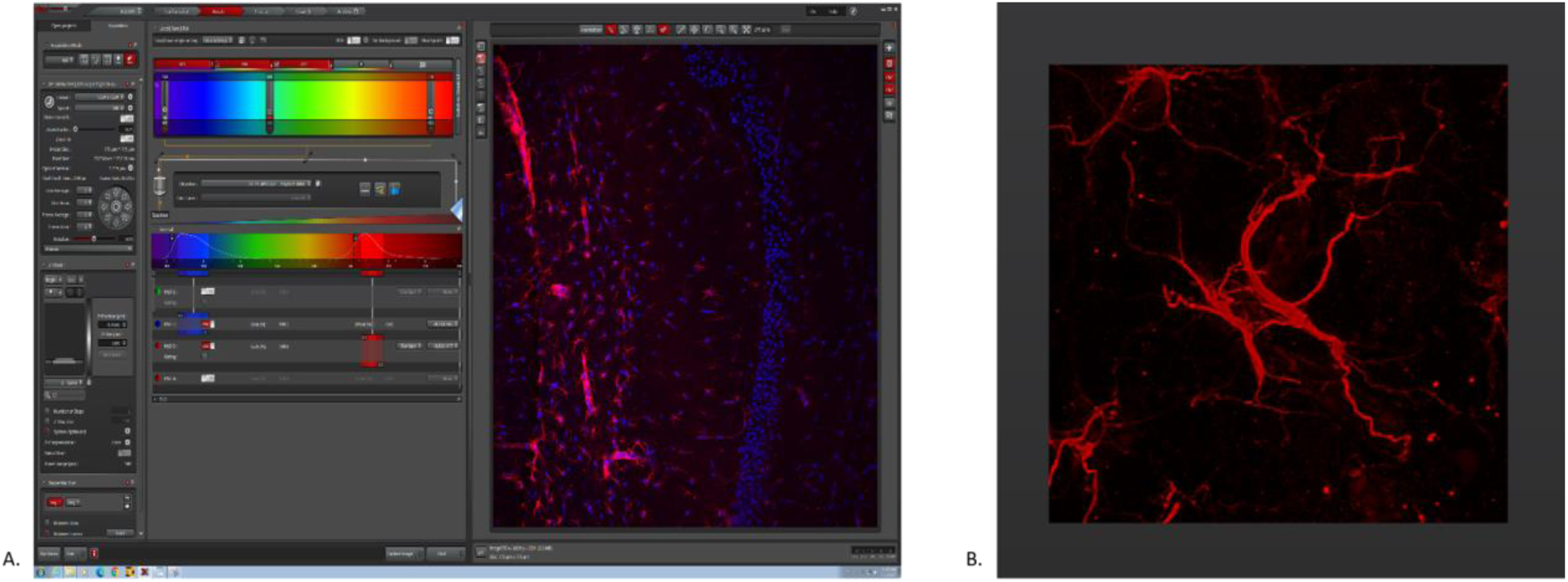
Representation of acquisition settings used in the Leica Confocal SP8 system. A. Parameters set during the acquisition of 20X magnification focusing on the CA1 area stained with GFAP (Red) and DAPI (Blue) B. High resolution of individual astrocyte (63X magnification) picked from the stratum radiatum layer.

**NOTE**: The step size to be chosen greatly depends on what analysis, the researcher wants to perform with the astrocytes. For instance, if someone is looking to characterize the structural analysis or the branching complexity of the astrocyte we would recommend having a smaller step size to have a detailed structural outlook of the astrocytes.

32. Click on “Start Experiment” to start taking acquisition of all the three channels. For an individual astrocyte this could take between 5-8 mins depending on the step size and the volume of astrocyte under consideration.
33. Deconvolution: To obtain a crisp image and to reduce the signal to noise (S/N) ratio we recommend performing deconvolution of the images. The lightening function from the Leica Confocal SP8 allows one to deconvolute the images before they are exported to imaris for further analysis. The parameters to be taken into account are regularization, iteration and smoothening. These parameters can be adjusted according to the image quality and the type of staining. The settings used for our image acquisition are found in Figure 6 A, B.

**Figure 6:**
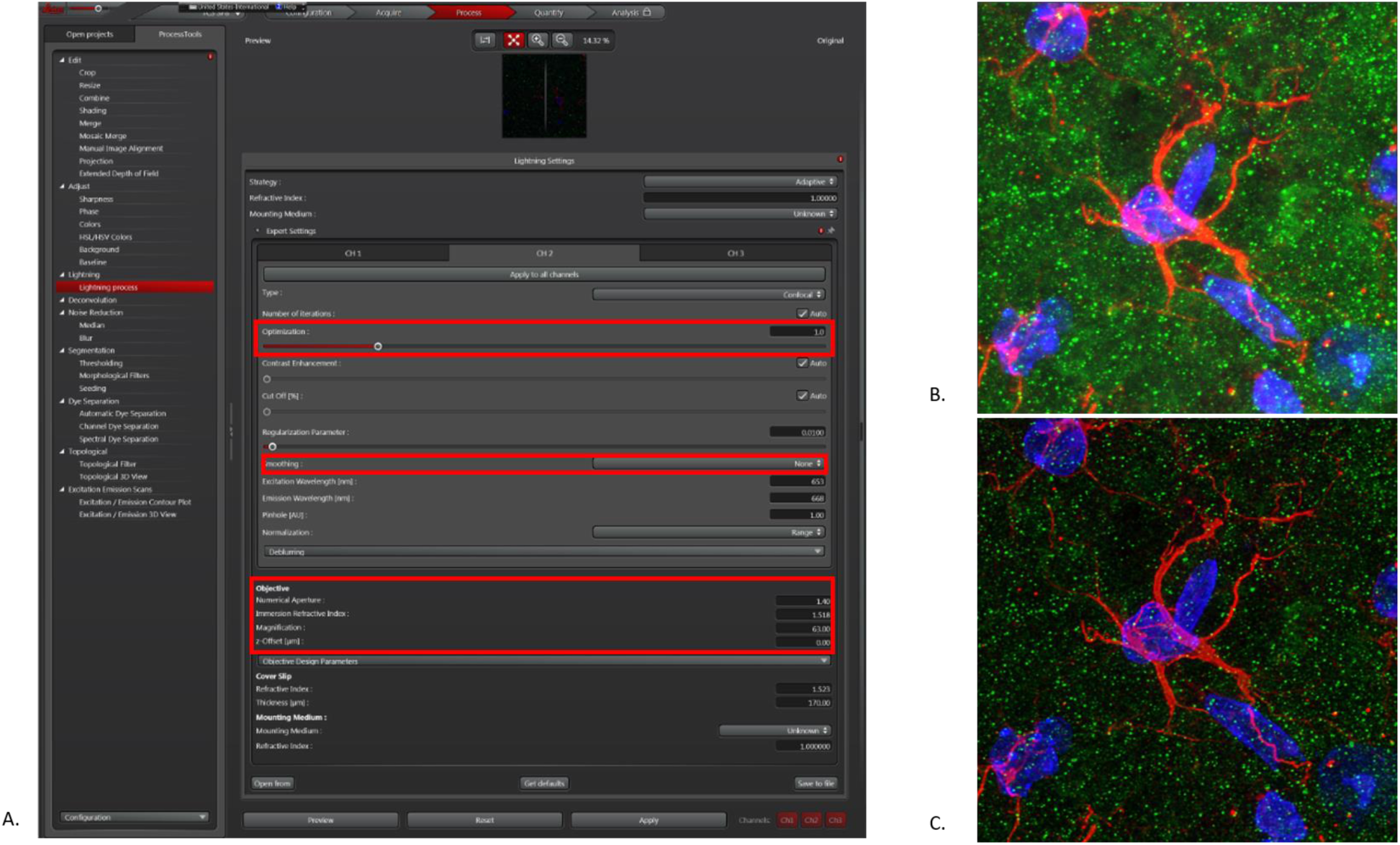
Deconvolution using the lightening function of the images acquired. A. Deconvolution parameters set during the acquisition of the GFAP channel. Of importance are the parameters marked using the red boxes. Switch off the smoothening function for the GFAP channel. B. 63X magnification astrocytes stained with DAPI (Blue) and C1q (Green) before deconvolution. C. After deconvolution images of the astrocyte. Deconvoluting the image highly reduces the background signal and improves the signal to noise ratio.

#### CRITICAL

Switch off the smoothening function for the GFAP channel, which might cause changes in the astrocyte structure and therefore affect the morphometric analysis.

#### CRITICAL

The regularization parameter is paramount to having a better signal to noise ratio. It represents to what extent a signal is interpreted as background or noise. Having an accurate adjustment of this parameter is essential to avoid false positives.

**NOTE**: If the confocal doesn’t have an inbuilt deconvolution software, one can use the deconvolution function from the Imaris Package.

34. Files can be saved in the appropriate “xxx.lif” format and can be exported to Imaris directly for quantification of synaptic pruning.

### Digital Processing: Measurement of Synaptic Engulfment

#### Timing: 15 min per astrocyte

This part is essential for the measurement of the synaptic engulfment of the C1q protein by the astrocytes. The procedure involves 3D volumetric reconstruction of individual astrocytes and colocalization analysis to calculate the C1q protein inside the entire volume of the astrocyte. PSD95, post synaptic marker has also been used as a reference to calculate the synaptic engulfment. Correct rendering of astrocyte volume forms a major step in the further colocalization analysis.

35. Since the “xxx.lif” format cannot be directly exported to Imaris, it can directly be converted to “xxx.ims” format in the Imaris converter.

#### CRITICAL

For a ubiquitously expressed protein, it might be necessary to perform a background subtraction, with a filter width of 1 μm. This improves the signal to noise ratio. The background subtraction can be done in the imaris package by clicking on the image processing and selecting background subtraction for the respective channel. In our case, the deconvolution using the lightening function from the Leica Confocal SP8 was sufficient to obtain high resolution imaging for all the channels.

36. Open the “xxx.ims” file in the Imaris software. Select “Add new surface” (blue oval like structure) from the tab menu.
37. The creation of the surface around the astrocyte takes place in 4 steps. In order to segment individual astrocyte, select “Segment only a region of Interest” under the “Algorithm Settings” box. Select the region of interest by dragging in through the X-Y plane. One can use pull arrows to restrict the region of interest (1/4).
38. In the next step, unselect the smooth option for the astrocyte. The smooth function as described before can affect the morphology of the astrocytes and thereby interfering with the final analysis (2/4). An overall visual guide to surface construction around the astrocyte is provided in Figure 7 A.
39. Next, we render the surface of the astrocyte by adjusting the signal threshold corresponding to the GFAP channel. One can manually slide the bar, under the “Threshold” tab to select the surface. Although there is an automatic threshold which is suggested by the Imaris, we suggest using the manual way to set the threshold in a way that the surface covers the complete astrocyte (3/4) (Figure 7 B).

**Figure 7:**
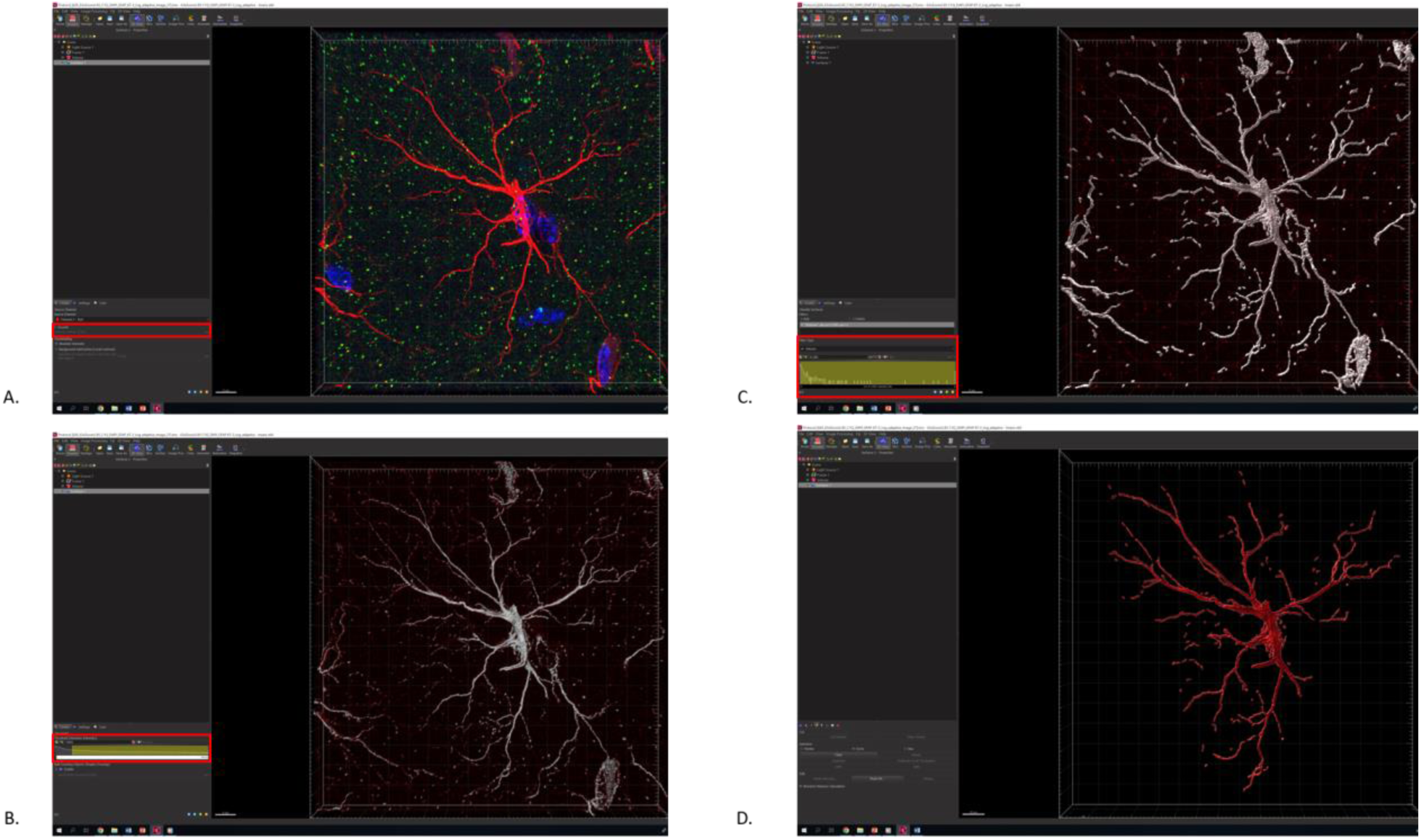
A visual guide through the four different steps of surface building on the astrocyte. A. In the step 2/4 select the red channel (GFAP), disable the smoothening option to avoid changes to the morphology of astrocytic surface B. In the step 3/4 the intensity threshold is selected for the red channel (For a detailed understanding of how to balance the threshold refer to Figure 8). While Imaris provides an automatic threshold for the surface reconstruction, we recommend to adjust the threshold manually for a more consistent and better rendering of the astrocyte. C. While the surface construction of the astrocyte, there are small unwanted disjoint signals produced in the background. In the step 4/4, we filter these small disjoint signals using a volume threshold filter D. The surface of the astrocyte is rendered.

#### CRITICAL

This step is crucial and plays an essential role in directly affecting the colocalization analysis. Correct rendering of astrocyte is essential to quantify the synaptic engulfment. Both over rendering and under rendering can provide false sets of data. We therefore recommend to do it on a trial and observe basis, once a surface is created, match with the original image, and see if the surface is rendered in the proper way (Figure 8). If not, come back to step 39 and repeat it again, until a proper rendering is achieved!

40. In the next step we can adjust the additional rendering required for achieving the volumetric rendering of astrocyte. For this step, set the filter to volume and manually adjust the threshold to remove small particle. We used a volume filter of 0.2 um^3^, to exclude unwanted particles. However, if the experimenter feels that additional small disjoint particles need to be removed, this parameter can be changed accordingly. However, to avoid any kind of bias leave the same setting for all the astrocytes (4/4) (Figure 7 C).
41. Click the green arrow button on step 4 to complete the rendering of the astrocyte. If desired rendering is not achieved, the user is advised to start again from step 39. This loop is to be followed until the desired rendering of astrocyte is achieved (Figure 7 D).

**Figure 8:**
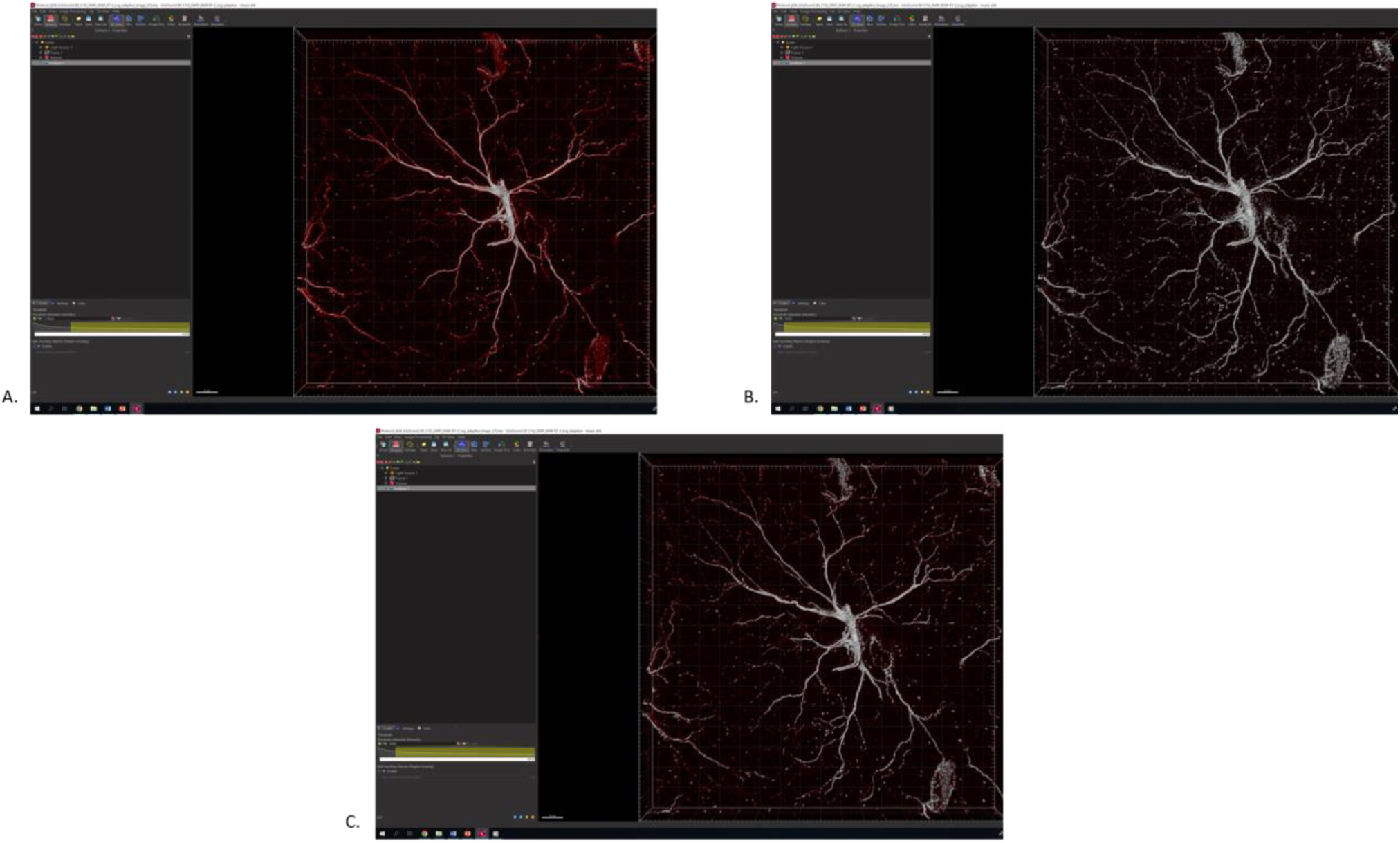
Creating the balance for a correct threshold for 3D rendering of astrocyte. A. Example of under representation of the surface of the astrocyte. In this case, the threshold is not adequately selected and several astrocytic filaments are not covered by the surface function. B. Over rendering of the surface. Having the threshold set too high, would introduce unnecessary and unreliable processes on the astrocyte. C. A balanced threshold should always be selected for the correct rendering of the astrocyte. The rendered astrocyte are now processed to further remove any other background signals.

**NOTE**: If a user intends to perform morphometric analysis of the astrocytes along with the colocalization study, Imaris provides an option to export all the statistics regarding the structural complexity and the branching details of the astrocyte. Click on the “Statistics” tab. Select all the parameters that you intend to study in the dropdown list “Export Statistics on Tab Display to file”. If the user intends to study all the parameters, select “Export all statistics to file”. The file gets automatically saved in the “xxx.csv” format. These sets of data can be analyzed further.

42. The next step is to process the rendered astrocyte (Figure 9 A). Usually even after careful surface reconstruction of the astrocyte, there are several unwanted signals or sometimes processes from other astrocytes which need to be filtered out before proceeding to the next step. This is achieved by selecting all the unwanted background signal (Figure 9 B) using the “Circle Selection Mode” (Figure 9 D) and then choosing the delete option under “Edit” tab (Figure 9 E). These unwanted signals are now filtered from the astrocyte (Figure 9 C).
43. The next step is to mask the surface of the astrocyte. To achieve this, click on the “Edit” tab under a newly built surface. Click “Mask All” option (Figure 10 A, B).
44. The mask channel box opens. Under the channels select GFAP (Red) channel. Set voxels outside the surface to 0. This is to eliminate any signal which is outside the surface that has been rendered. This step further reduces any false positives resulting from the background signal (Figure 10 C, D).

**Figure 9:**
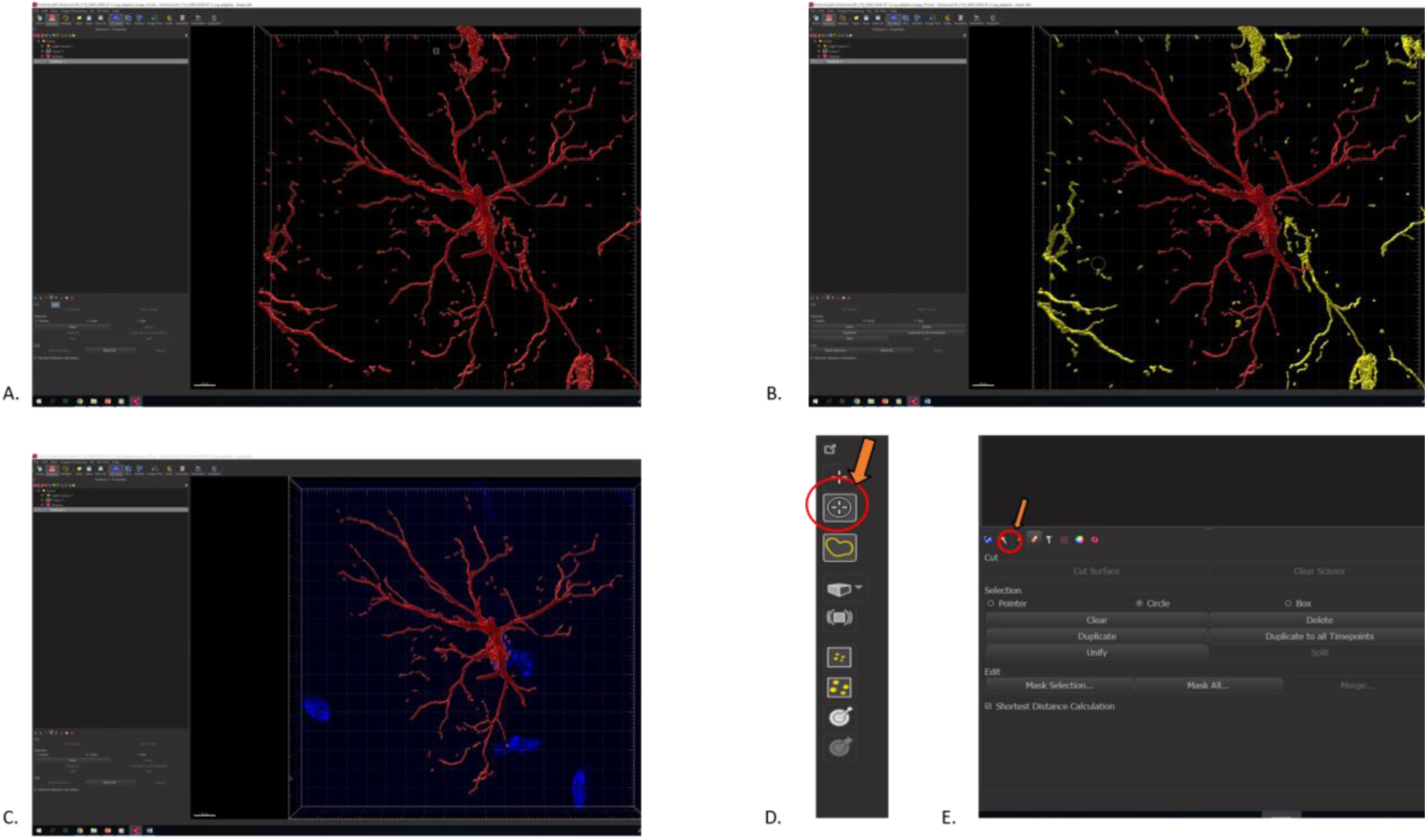
Processing of rendered astrocyte. A. The rendered astrocyte produced after step 4/4 of surface creation. There are several other background signals as well as processes from other astrocytes. These need to be filtered out before performing the colocalization analysis. B. The unwanted signals and processes are selected and are represented in yellow C. Processed astrocytes with DAPI staining to determine and match the processes with the respective astrocyte. D. The unwanted signals are selected by using the “Circle Selection Mode” shown by orange arrow in the upper right corner. E After selecting all the unwanted background signal we select the edit option (pencil like icon on the lower left side indicated by the orange arrow) and click on the delete option.

**Figure 10:**
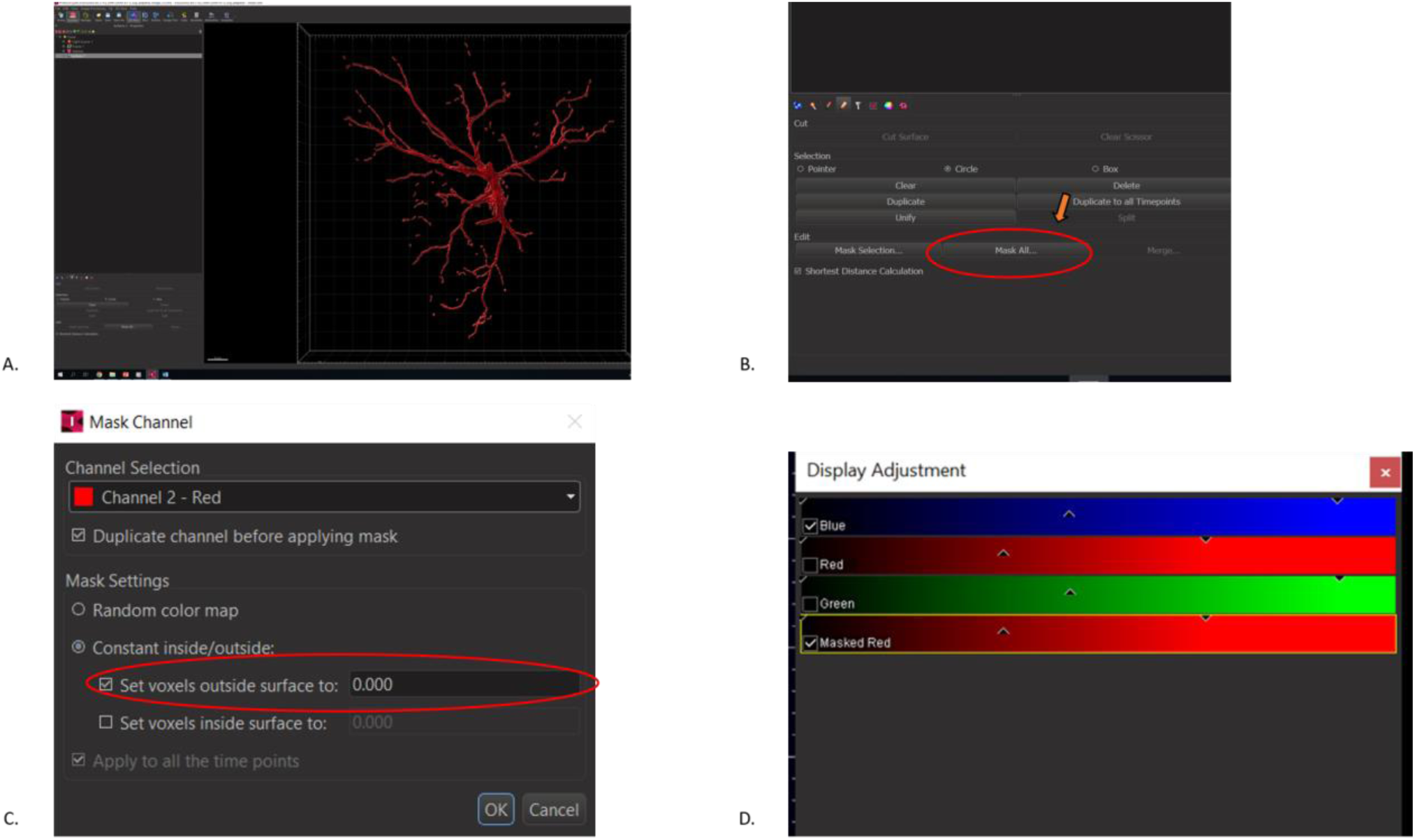
A visual guide to mask the surface of the astrocyte. A. Rendered and processed astrocyte B. On the edit option (pencil like icon), select “Mask All” option (Shown by the arrow) C. A drop down selection box appears. Select the channel in which the astrocyte is present (red), and set the voxels outside surface to 0. D. The masked surface of the astrocyte appears and in the display adjustment box, in addition to blue, red and green channel, the masked red channel appears.

**NOTE**: To quantify the synaptic engulfment, we use the colocalization analysis. The coloc analysis of Imaris analyzes the contact between the C1q protein (Green) and the astrocyte surface. Using the masked red channel we can calculate the contact between the C1q and the astrocyte. The percentage of region of interest (Astrocyte) colocalised with the neuronal tag can be exported in a separate file. In this part we are focusing on the colocalization between C1q and GFAP (Astrocytes). However this analysis can be extended to different post synaptic and pre synaptic markers.

45. Start the Coloc tab in the Imaris (Figure 11 A).
46. Imaris allows the selection of 2 channels at once which are to be analyzed for the colocalization. In the first channel (A) select green (C1q) (Figure 11 B). The next step is to set the threshold intensity for the green channel. While Imaris provides an automatic way to determine threshold, we recommend not to use it because of inconsistent and unreliable results. The better way to analyze the threshold for green channel is by setting a manual threshold. This can be achieved by randomly selecting 10 green dots and averaging the intensity to set the threshold for the green channel. Left click and drag on a particular C1q protein in a way that the boundary set by Imaris completely covers the protein (Figure 11 C). Find the mean value of the 10 randomly selected C1q protein signals and set the threshold intensity of channel A.

**Figure 11:**
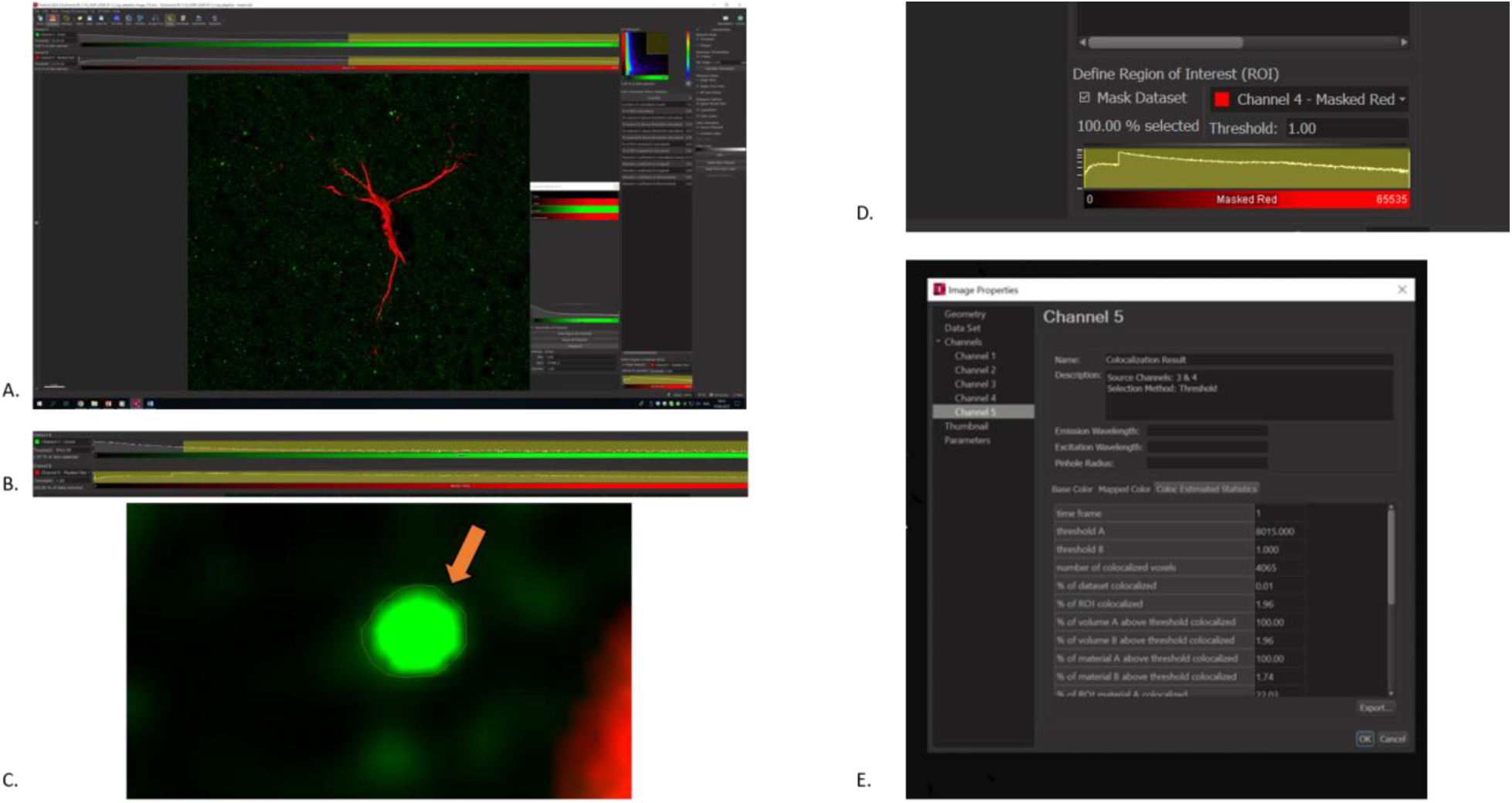
Guide for the colocalization analysis with Imaris. A. Open the Coloc tab which is present at the side of “Image Processing” option. B. The intensity threshold for channel A (green) is selected by averaging the intensities of 10 unambiguous signal selected by setting the boundary around each signal. For the channel B (masked red), the threshold is set at 1. C. Correct way of setting the boundary around the positive green signal. Make sure that the yellow boundary covers all of the individual green signal D. In the “Define Region of Interest (ROI)” box in the lower right corner, select mask dataset and select the masked red channel in the channel box E. The colocalization analysis produces a set of data which define % of colocalization. These values can be exported to a separate excel file. The % of ROI colocalised is of interest to us as it gives us the measurement of synaptic engulfment by the astrocyte.

#### CRITICAL

While Imaris recommends to set the threshold for both the channel using the 2D histogram, in our experience it is better to proceed with manual threshold instead of using the 2D histogram, as it produces inconsistent results.

#### CRITICAL

While encircling the boundary of a positive C1q signal, care should be taken to completely encircle the boundary. Wrong acquirement, could produce falsified results in the colocalization analysis.

47. In the channel B select the masked GFAP and set the intensity threshold for the channel B (Figure 11 B). This is relatively easy as compared to the setting of the threshold for channel A. During the masking protocol we eliminated the background signal by threshold masking of the original GFAP channel. Therefore, the threshold intensity of this channel can be directly set at 1.
48. Towards the bottom in the right side, select the define “Region of Interest” tab. This is a crucial step in the analysis of the final set of the data. Since we intend to calculate the colocalization over the whole volume of astrocyte, we select the masked GFAP channel and set the threshold as 1 (Figure 11 D).
49. The last step of the analysis is to “Build Coloc Channel”. A new tab opens with different parameters analyzed. The whole file can be exported as a “xxx.csv” file. To our interest, is the % of ROI colocalised. This gives us an estimation of the amount of C1q tag present in the whole volume of the astrocyte. This can be transferred to a master excel file consisting of data from several other astrocytes and in both control and treatment samples for the final statistical analysis (Figure 11 E).

## Expected outcomes

In this paper, we present a fast and efficient method to characterize the synaptic engulfment of neuronal tag in an *in vitro* slice culture treatment of amyloid beta oligomers by the astrocyte (Figure 12, 13). The colocalization analysis protocol serves as an indirect measure for the synaptic pruning process. Amyloid beta oligomers are known to negatively affect the spine density and synapses. C1q, a complementary protein tag, serves as a destruction signal for the neurons to be engulfed by the glial cells. The CA1 area of the hippocampus plays an essential role in memory and learning and inducing of long term potentiation (LTP). From our previous research, the treatment of hippocampal slice with 50nM Aβ_42_ causes a significant decline in the CA1-LTP (Rammes *et al*., 2018). This protocol was developed to understand the molecular ongoing particularly in the astrocyte when the slices are incubated with Aβ_42_. The % ROI colocalization gives the measurement of the amount of synaptic tag engulfed by the whole volume of the astrocytes.

**Figure 12:**
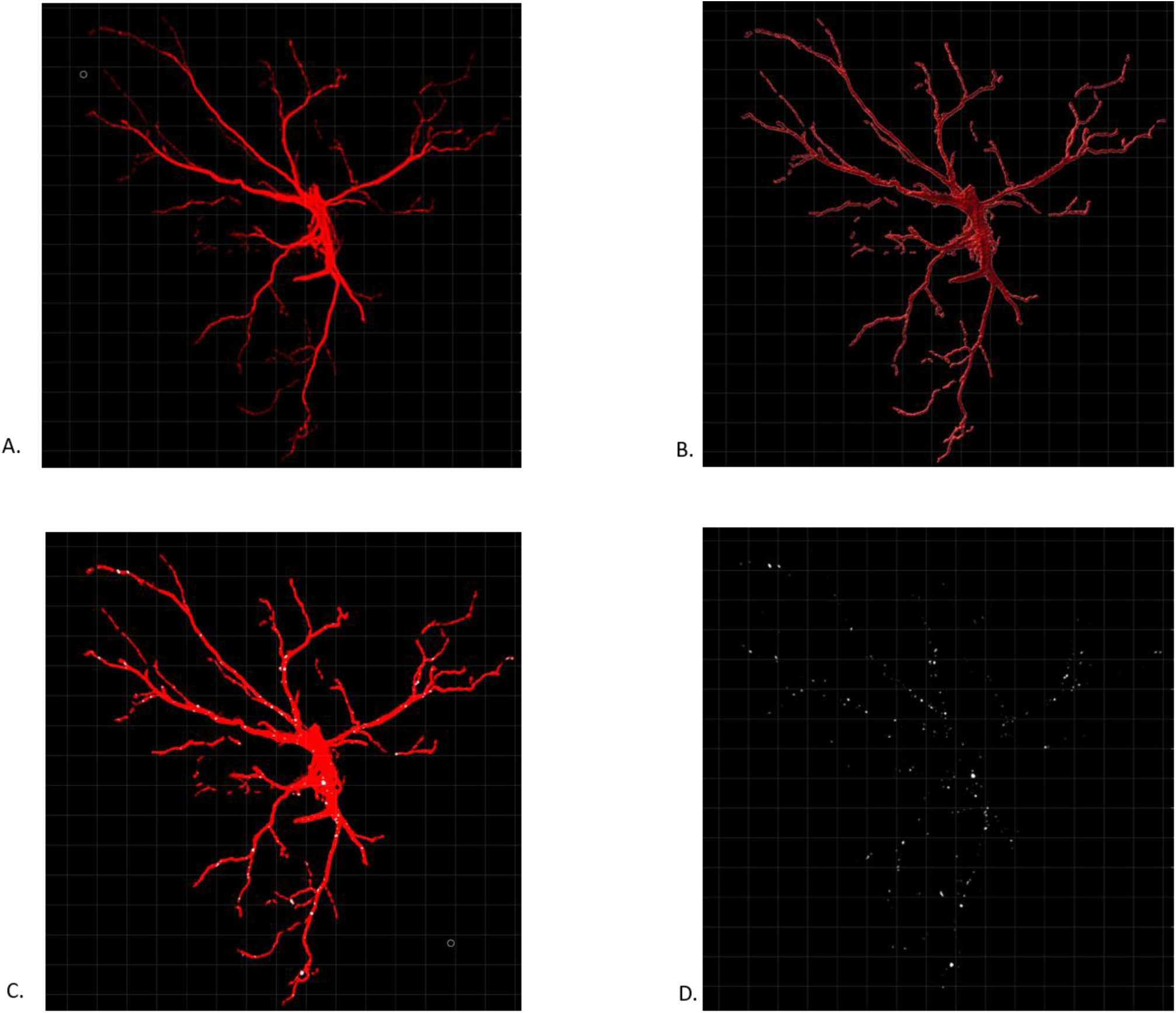
Volumetric rendering and synaptic engulfment of the astrocyte. A. Original processed image of the astrocyte B. Volumetric 3D rendered image C. Astrocyte (Red) with colocalization of C1q in white D. Colocalization shown in white

**Figure 13:**
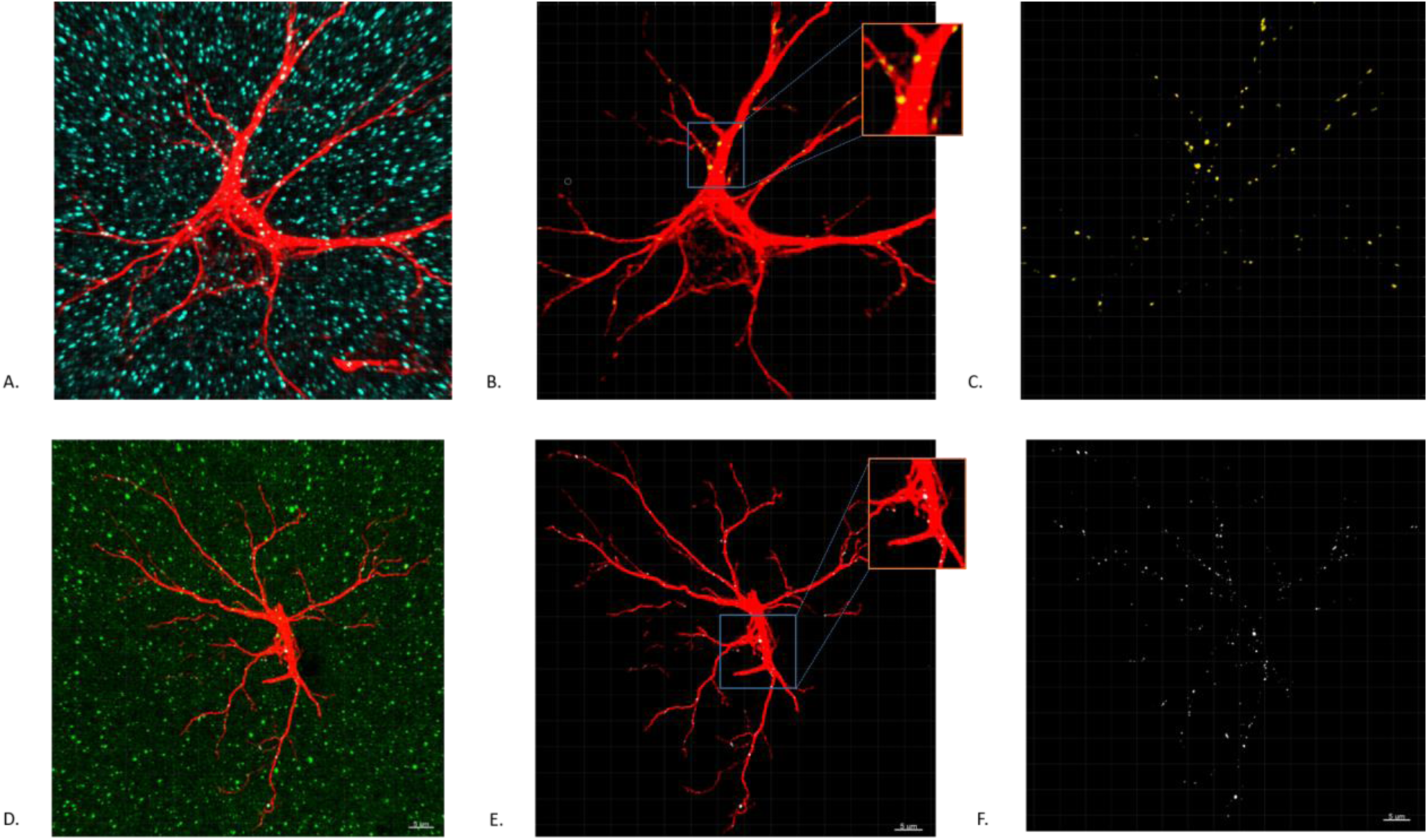
Synaptic engulfment of post synaptic marker PSD95 and complementary tag C1q. A. Individual high resolution astrocyte (shown in red) and post synaptic marker PSD95 (shown in light blue) B. Processed astrocyte showing colocalization with PSD95. The colocalization points are shown in yellow inside the astrocyte. C. The yellow points represents the colocalization between astrocyte and PSD95 D. Astrocyte (shown in red) and the “eat-me signal” C1q protein (shown in green) E. Processed astrocyte (Red) with C1q colocalization shown in white F. The colocalised points between astrocyte and C1q are represented in white.

## Quantification and statistical analysis

In order to have a meaningful number of samples to test the significance, we recommend having at least 6 mice per group. It is advised to average the data from at least 15-20 astrocytes (from every mice brain) to have a well-represented data set. The statistical analysis can be performed by common programs such as Graph Pad Prism or SPSS. The statistical test to be used for analysis, greatly depends on the type of experiment and number of groups. If the control group is to be compared with an experimental group only, it would be advisable to use the two-tailed unpaired T-test.

## Limitations

The protocol described here and the rationale behind utilizing this protocol applies without any bias to animals of age group 6-8 weeks. Older animals, however, due to the age parameter have more reactive astrocytes as compared to the younger ones. Therefore, the reactive nature of the astrocyte whether due to the treatment with amyloid beta oligomers or due to the age parameter cannot be said with a certainty. The quality of the tissue and the protocol used for collecting the samples greatly impacts the quality of rendering as well as the colocalization study. The protocol described in this paper best applies to free floating slices. If the same is applied to the frozen slices, the rendering would be inaccurate. The quantification of intensity of the fluorophores is a relative parameter, and it should always be normalized with respect to the negative control, to obtain a reliable analysis of colocalization.

## Troubleshooting

### Problem 1

Poor GFAP signals/astrocytic processes are unclear

### Potential solution

The protocol described in this paper suits best for the free floating slices. We have observed that when the same protocol is repeated for the frozen section, the astrocytic processes are not clear and the branching complexity usually cannot be inferred from such data sets. We do have a separate protocol for accessing the colocalization in the frozen section. Briefly, instead of using PFA as a fixative agent we use chilled acetone: isopropanol (1:1) as a fixative agent. The fixation is done by inserting the slides containing brain slices inside the acetone: isopropanol solution for about 25 min. This is followed by the permeabilization by 0.3% triton-X in PBS for a span of 20 minutes followed by washing with PBS three times with 1X PBS and then blocking with 10% NGS at room temperature for an hour. This is followed by antibody incubation as described in the current protocol.

### Problem 2

Poor PSD95 signal obtained from the confocal microscope

### Potential solution

Initially we started out with incubation of primary antibody of PSD95 for overnight at 4ºC, however the signal obtained was sparse and unclear. We recommend therefore incubating the primary antibody for PSD95 for 2 days.

### Problem 3

The brain slice breaks while cutting in the cryotome/ there are cracks in the slices collected

### Potential solution

The brain slice if not kept flat, will result in unequal thickness of slices. This might result in breaking of slices or having folds in the slices. In such cases there are chances that obtaining the complete hippocampus becomes difficult. An easier way is to put the slices horizontally on the solidified OCT mounting medium and softly pressing it by using a brush. The slice (30 μm) needs to be flat for efficient penetration of antibodies. However, if the slices are flat and there are still cracks in the slices, consider changing the object temperature and knife temperature.

### Problem 4

While 3D reconstruction of the astrocyte, there are small unnecessary signals which are far from the main astrocyte under construction.

### Potential solution

There is a twofold way to solve this issue. In the step 4 of the volumetric rendering of the astrocyte, extra signals outside the main astrocyte can be controlled by setting a volume filter as described in step 40. This can be used on a trial and observe basis to exclude the unnecessary background signals. However, if there are still unspecific signals after the 3D rendering of the astrocyte, they can be removed manually by selecting “Circle Selection Mode” on the right side of the Imaris panel. Select the background signal by clicking CTRL and selecting the signals using the right cursors. After all the unnecessary signals are selected, they can be deleted by clicking “pencil” like icon (Edit) on the lower left side and then selecting the delete option.

### Problem 5

There is ambiguity regarding the processes whether from a single astrocyte or more than 1 astrocytes.

### Potential solution

There is always a possibility that there are processes from other astrocytes in the same frame. The easiest way to detect or possibly have a clue whether the processes belong to the same astrocyte, is to have a DAPI (nuclear) staining. If the processes from other astrocytes are detected on the same frame as the astrocyte under the consideration, they need to be manually removed as suggested in the solution to Problem 4.

### Problem 6

The staining of astrocyte and synaptic marker is not clear.

### Potential solution

While there could be a lot of parameters impacting the staining, starting from the incubation of slice to the immunofluorescence protocol. We particularly advise to be careful with 2 parameters, if the immunofluorescence protocol was performed correctly. First, the incubation period of slices with the amyloid beta peptide need to be correctly monitored. If the slices are overlapping or there are bubbles in the chamber holding the slice, the health of the slices are greatly hampered. Second, the sections obtained from the cryotome should be flat and without any cracks.

## Resource availability

### Lead contact

Further information and requests for resources and reagents should be directed to and will be fulfilled by the lead contact, Gerhard Rammes.

### Materials availability

The study did not generate any new reagents.

## Data and code availability

The study did not generate/analyze new code.

## Acknowledgments

The work was supported by DFG research grant (RA 689/12-1) and funding provided by the China Scholarship Council (CSC) (File No. 202008080068). We would like to thank Dr. Ritu Mishra (Core

Facility, Translatum) at Klinikum Rechts der Isar for access to the confocal microscopy facility and the Imaris analysis system. We would also like to thank Mr. Andreas Blaschke for helping us with the amyloid beta preparation.

## Author contributions

AKP and QS developed the protocol. AKP wrote the manuscript. AKP, KT and QS performed the experiment. KT and QS provided the images for the manuscript. GR supervised the whole work.

## References

Allen, N. J. and Eroglu, C. (2017) ‘Cell biology of astrocyte-synapse interactions’, Neuron, 96(3), pp. 697–708.

Chung, W.-S., Allen, N. J. and Eroglu, C. (2015) ‘Astrocytes control synapse formation, function, and elimination’, Cold Spring Harbor perspectives in biology, 7(9), p. a020370.

Garland, E. F., Hartnell, I. J. and Boche, D. (2022) ‘Microglia and Astrocyte Function and Communication: What Do We Know in Humans?’, Frontiers in Neuroscience, 16.

Györffy, B. A. et al. (2018) ‘Local apoptotic-like mechanisms underlie complement-mediated synaptic pruning’, Proceedings of the National Academy of Sciences, 115(24), pp. 6303–6308.

Hong, S. et al. (2016) ‘Complement and microglia mediate early synapse loss in Alzheimer mouse models’, Science, 352(6286), pp. 712–716.

Iram, T. et al. (2016) ‘Megf10 is a receptor for C1Q that mediates clearance of apoptotic cells by astrocytes’, Journal of Neuroscience, 36(19), pp. 5185–5192.

Kovács, R. Á. et al. (2021) ‘Identification of neuronal pentraxins as synaptic binding partners of C1q and the involvement of NP1 in synaptic pruning in adult mice’, Frontiers in immunology, 11, p. 3792.

Li, S. et al. (2011) ‘Soluble Aβ oligomers inhibit long-term potentiation through a mechanism involving excessive activation of extrasynaptic NR2B-containing NMDA receptors’, Journal of Neuroscience, 31(18), pp. 6627–6638.

Perez-Catalan, N. A., Doe, C. Q. and Ackerman, S. D. (2021) ‘The role of astrocyte-mediated plasticity in neural circuit development and function’, Neural Development, 16(1), pp. 1–14.

Rammes, G. et al. (2018) ‘The NMDA receptor antagonist Radiprodil reverses the synaptotoxic effects of different amyloid-beta (Aβ) species on long-term potentiation (LTP)’, Neuropharmacology, 140, pp. 184–192.

Tau, G. Z. and Peterson, B. S. (2010) ‘Normal development of brain circuits’, Neuropsychopharmacology, 35(1), pp. 147–168.

Tierney, A. L. and Nelson III, C. A. (2009) ‘Brain development and the role of experience in the early years’, Zero to three, 30(2), p. 9.

